# Construction and Integration of Three *De Novo* Japanese Human Genome Assemblies toward a Population-Specific Reference

**DOI:** 10.1101/861658

**Authors:** Jun Takayama, Shu Tadaka, Kenji Yano, Fumiki Katsuoka, Chinatsu Gocho, Takamitsu Funayama, Satoshi Makino, Yasunobu Okamura, Atsuo Kikuchi, Junko Kawashima, Akihito Otsuki, Jun Yasuda, Shigeo Kure, Kengo Kinoshita, Masayuki Yamamoto, Gen Tamiya

## Abstract

The complete sequence of the human genome is used as a reference for next-generation sequencing analyses. However, some ethnic ancestries are under-represented in the international human reference genome (e.g., GRCh37), especially Asian populations, due to a strong bias toward European and African ancestries in a single mosaic haploid genome consisting chiefly of a single donor. Here, we performed *de novo* assembly of the genomes from three Japanese male individuals using >100× PacBio long reads and Bionano optical maps per sample. We integrated the genomes using the major allele for consensus, and anchored the scaffolds using sequence-tagged site markers from conventional genetic and radiation hybrid maps to reconstruct each chromosome sequence. The resulting genome sequence, designated JG1, is highly contiguous, accurate, and carries the major allele in the majority of single nucleotide variant sites for a Japanese population. We adopted JG1 as the reference for confirmatory exome re-analyses of seven Japanese families with rare diseases and found that re-analysis using JG1 reduced false-positive variant calls versus GRCh37 while retaining disease-causing variants. These results suggest that integrating multiple genome assemblies from a single ethnic population can aid next-generation sequencing analyses of individuals originated from the population.

## INTRODUCTION

The complete human genome sequence^1,2^ has been an invaluable resource for both basic research in human genetics and clinical diagnosis. The complete genome sequence is currently used as a reference for mapping the enormous number of short reads generated using major next-generation sequencing (NGS) techniques^3,4^, is thus also called “the reference genome”. Because the short reads generated in NGS studies are approximately 100–300 bp in length, mapping them to the reference genome is an indispensable step for calling single nucleotide variants (SNVs) and short insertions and deletions (indels) in the sample individuals. The coordinate system of the reference genome is used for biological and medical annotations, such as the position or sequence of specific genes, or sites of causal variants associated with both rare and common diseases. Therefore, the reference genome is one of the most foundational resources in human genetics, and as such, it is maintained and continually updated by the Genome Reference Consortium (GRC). The latest and second-latest versions of the reference genome (GRCh38/hg38 and GRCh37/hg19, published in 2013 and in 2009, respectively) are nearly complete, and both are widely used for NGS analyses and genome annotations^5,6^.

The reference genome was constructed using a hierarchical shotgun sequencing strategy in which fragmented genomic DNA segments cloned in bacterial (BAC) or P1-derived (PAC) artificial chromosome libraries are arranged into a correct physical map to guarantee that the reference genome was haploid (mosaic)^1^. The assembled contigs or scaffolds were then anchored on each chromosome using information from genetic and radiation hybrid (RH) maps, which have thousands to tens of thousands of sequence-tagged site (STS) markers in linkage groups (i.e. chromosomes). It should be noted that these genetic and RH maps are original information sources used to construct the reference genome and not derived from the reference genome itself.

Although the reference genome is a resource of unparalleled value, several of its characteristics are not ideal for application to NGS analyses, particularly for some populations^7^. For example, although the reference genome is constructed using genetic information from multiple donors, each clone comprising the resulting reference genome is derived from either haploid genome of a particular individual. As such, the reference genome inevitably harbors rare or even private variants. Over 90,000 rare variants were used as a reference allele including disease-susceptibility variants for thrombophilia and type 2 diabetes^8,9^. Inclusion of such variants in the reference can lead to erroneous and confusing results of short read mapping or variant calling^9^. As the NGS analyses typically assume that the reference allele is the ancestral, healthy, or major allele for any variable site, the inclusion of such rare alleles may also confuse subsequent interpretations.

Another possible problem associated with the reference genome is that the samples used for its construction are biased toward African and European ancestries. For example, >70% of the reference genome is composed of a BAC library known as RP-11 (aliased RPCI-11)^1^ from a donor with both African and European ancestry^10^. With the exception of one donor with an Asian background, all of the donors had a European background resulting in the composition of an Asian haplotype for 4.3% of the reference genome^1,10^. In addition, one recent study revealed a lack of population-specific sequences in the reference genome^11^, whereas another discovered thousands of structural variants (SVs) in world-wide samples^12^. These issues can also complicate short-read mapping and variant callings.

Several studies have examined ways to overcome the above-mentioned drawbacks to the reference genome. Dewey et al^13^ proposed modifying the reference genome by substituting its minor variants with the major variants from African, Asian, or European populations^13^. The resulting modified reference genome was better-suited for genome analyses of sample individuals with matched population backgrounds. Several studies^14–17^ utilized a genome graph, which is an extended reference genome represented as a graph harboring known variants. Other studies have proposed the addition of sequences not included in the reference genome^11,12,18,19^. However, these proposed adjustments are based largely on variants discovered using the reference genome itself, albeit only partially, in a circular fashion, some reference bias could remain.

One promising approach to address these problems is to construct new reference genomes specific to ethnic populations of interest^20^. Although costly, highly contiguous *de novo* assembly—independent reconstruction—of the entire human genome is now feasible using, for example, Pacific Biosciences (PacBio) single molecule, real-time (SMRT) long reads (∼10 kb in length) and Bionano optical mapping, which generates a high-resolution physical map^18,21,22^. Combining of these approaches is known as ‘hybrid scaffolding,’ which is carried out in three steps: 1) PacBio long reads are *de novo* assembled to yield primary contigs; 2) Bionano raw data are also *de novo* assembled (independent of the PacBio assembly) to yield optical maps; and 3) the PacBio-derived contigs are scaffolded by the Bionano optical maps. This strategy is analogous to the hierarchical shotgun sequencing strategy used in the Human Genome Project^1^ with arrangements of long sequences from BAC/PAC on a physical map. Although assemblies generated in recent studies were highly contiguous and accurate, the assembled sequences were rarely anchored to a set of chromosomes, thus making their use as references for NGS analyses impractical. Moreover, a single haploid assembly from a single individual cannot be used to solve the rare reference allele problem. A notable exception is the KOREF genome sequence^20^, in which a Korean reference genome was constructed by *de novo* assembly of the genome sequence of a Korean individual, reconstructed as a set of chromosomes, and rare variants were substituted with short reads from 40 Korean individuals. However, the KOREF genome assembly was found to be less contiguous than long read-based assemblies because the primary sequencing platform was a short-read sequencer, and KOREF depended heavily on the reference genome because chromosome building was carried out by sequence-based alignment of scaffolds onto GRCh38.

Using a hybrid scaffolding strategy, in this study, we constructed a new reference genome, JG1, by integrating *de novo* assemblies of three Japanese individuals. After merging the three haploid assemblies constructed by hybrid scaffolding strategy, we defined major variants among the three (i.e., majority decision) and adopted them as the reference allele. We also positioned the scaffolds along chromosomes with the aid of conventional genetic and RH maps. We then assessed the extent to which JG1 represents the major variants in the Japanese population in terms of SNVs and SVs. As an example potential application, we also demonstrated the utility of using JG1 as a reference genome in NGS analyses aimed at identifying the causal variants of several rare diseases.

## RESULTS

### Construction of JG1

To construct a genome sequence with population reference-quality, the population background of the reference genome should not significantly diverge from the backgrounds of sample individuals in order to reduce unnecessary variant calls that merely reflect the difference in the population background. In the case of our study, the donor should therefore be chosen from the Japanese population originating from the main island of Japan. In addition, we built the Japanese reference genome independent of the GRC reference genome in order to eliminate known ethnic biases toward African and European backgrounds as well as any other (and possibly unknown) biases. We therefore, performed *de novo* assembly of the Japanese human genome. Majority-based decision-making regarding multiple *de novo* assemblies was implemented as an effective way to avoid inclusion of rare reference alleles. This majority-based decision-making strategy produced a haploid genome sequence amenable to analyses using currently available and standard bioinformatics tools for NGS data.

We recruited three male Japanese volunteers, and they were given the sample names jg1a, jg1b, and jg1c (jg1a is the same individual as JPN00001^19^). Principal component analysis (PCA) based on the genotypes inferred by whole-genome sequencing indicated that the subjects were scattered within the cluster of the Japanese population (Figure 1a). G-banding analyses (Supplementary Fig. 1) indicated that all three individuals had a normal karyotype, although subject jg1a had a common pericentric inversion within chromosome 9, inv(9)(p12q13). Because it was difficult to assemble the pericentric region of chromosome 9 equally for all three subjects, this variation does not appear to have affected the assembly results (Supplementary Fig. 2).

**Figure 1.**
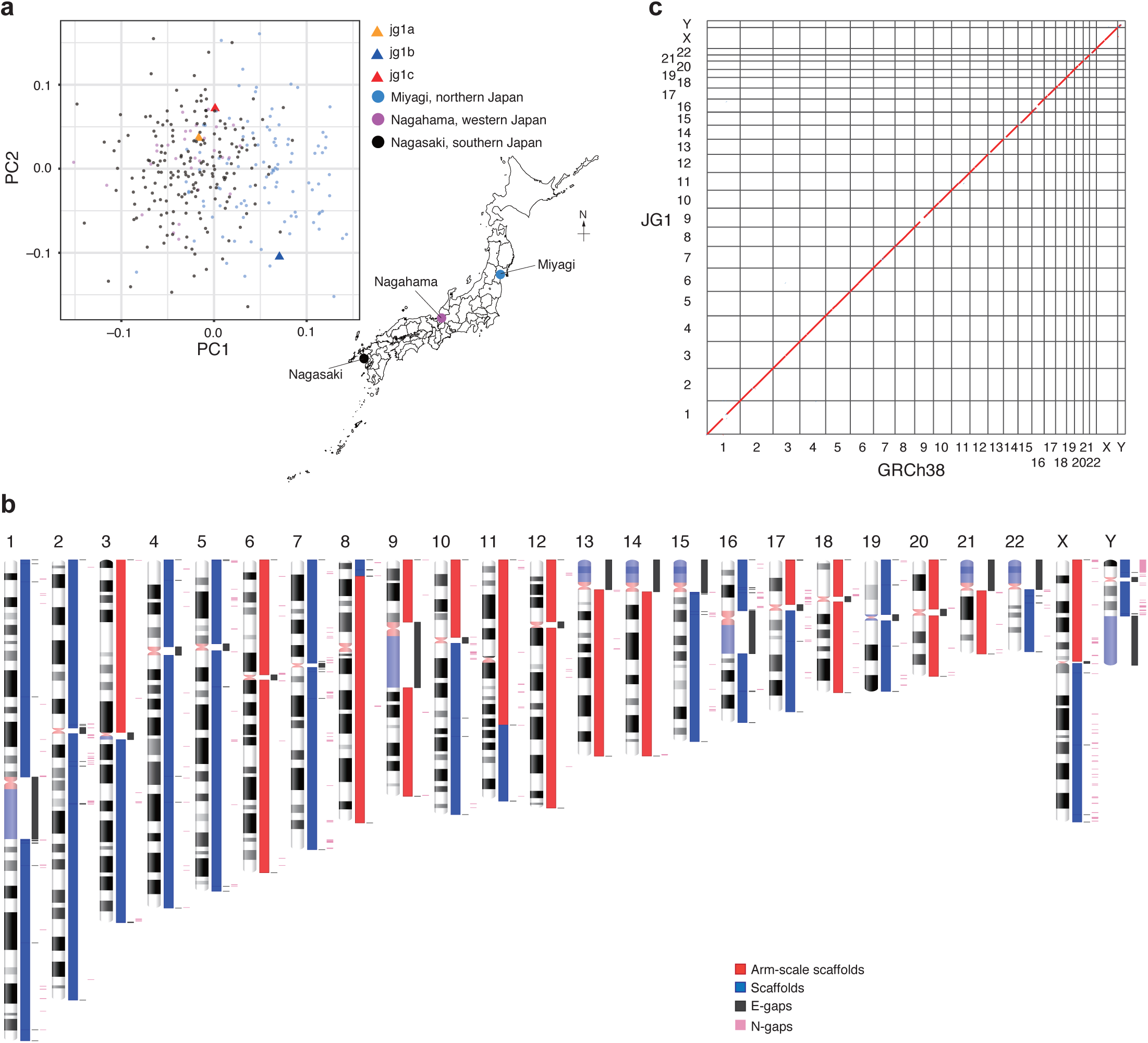
Construction of JG1. **(a)** PCA plot showing that the three sample donors are within the Japanese population cluster. **(b)** Idiogram showing the regions sequenced for each chromosome in JG1. Red and blue boxes indicate scaffolds; the red box spans an entire chromosomal arm. Dark gray boxes denote E-gaps, which represent links connected by genetic and RH maps or gaps inserted according to other evidence. Pink boxes denote N-gaps, which are unresolved regions linked by Bionano optical maps, or the putative PAR1 region in the Y chromosome. **(c)** Harr plot representing the co-linearity between the reference genome GRCh38 and JG1.

To construct a reference-quality haploid genome sequence, we integrated the three *de novo* assembled genomes (see Supplementary Fig. 3 for an overview; see Supplementary Tables 1-3 for materials). For each subject, we sequenced deeply (122× for jg1a, 123× for jg1b, and 128× for jg1c) using PacBio technology (Supplementary Fig. 4 and Supplementary Table 1) and then performed *de novo* assembly using Falcon software^23^. The *de novo* assemblies yielded 2,194, 2,227, and 2,120 primary contigs for jg1a, jg1b, and jg1c, respectively (Table 1). The contig N50 value was approximately 20 Mb for the three subjects (Table 1). Using ArrowGrid software^24^, the primary contigs were then error-corrected (polished) with the same long reads used for the initial *de novo* assembly.

**Table 1.** Assembly results.

|  | **Total length (bp)** | **Number of fragments** | **N50 (bp)** | **NG50 (bp)** | **Number of misassemblies** | **Reference** |
| --- | --- | --- | --- | --- | --- | --- |
| jg1a primary contigs | 2,855,392,439 | 2,194 | 20,631,146 | 19,227,791 | 1,912 | this study |
| jg1b primary contigs | 2,852,624,381 | 2,227 | 21,603,629 | 19,007,577 | 1,673 | this study |
| jg1c primary contigs | 2,851,554,649 | 2,120 | 19,616,169 | 17,539,317 | 1,673 | this study |
| jg1a scaffolds | 2,889,327,167 | 1,911 | 86,280,884 | 59,417,266 | 2,071 | this study |
| jg1b scaffolds | 2,880,572,022 | 1,893 | 59,380,744 | 57,047,228 | 1,762 | this study |
| jg1c scaffolds | 2,875,657,275 | 1,797 | 58,198,703 | 58,048,754 | 1,867 | this study |
| meta-scaffolds (jg1c + (jg1a + jg1b)) | 2,858,691,982 | 708 | 66,367,161 | 58,207,422 | 1,581 | this study |
| JG1 | 3,085,782,898 | 624 | 141,953,703 | 141,953,703 | 1,654 | this study |
| AK1 scaffolds | 2,904,207,228 | 2,832 | 44,846,623 | 39,609,866 | 2,138 | 21 |
| HX1 scaffolds | 2,934,082,568 | 5,323 | 21,979,250 | 20,700,129 | 2,688 | 22 |
| Swe1 scaffolds* | 3,127,010,000 | NA | 49,799,000 | NA | NA | 18 |
| Swe2 scaffolds* | 3,103,497,000 | NA | 45,443,000 | NA | NA | 18 |

\* Results of Swe1 and Swe2 scaffolds were obtained from Ameer et al<sup>18</sup>. All other results were calculated by using Quast-LG software<sup>60</sup> with the reference GRCh38 as the truth set.

We also obtained deep Bionano data for each subject (123× and 140× for two enzymes for jg1a; 160× and 175× for one enzyme for jg1b and jg1c, respectively; Supplementary Fig. 5 and Supplementary Table 2), and performed *de novo* assemblies of these data to generate optical maps (Supplementary Table 4). Each *de novo* assembly of the Bionano data was performed in two rounds (rough and full) to guarantee independence relative to the GRC reference genome (see Methods section). We then performed hybrid scaffolding between the PacBio-derived contigs and the Bionano-derived optical maps. The resulting hybrid scaffolds were then polished with 55×, 59×, and 57× Illumina short reads for subjects jg1a, jg1b, and jg1c, respectively (Supplementary Table 3). The number and N50 value of the resulting hybrid scaffolds were 1,911, 1,893, and 1,797, and 86.28 Mb, 59.38 Mb, and 58.20 Mb for subjects jg1a, jg1b, and jg1c, respectively. These and other assembly statistics were better than or comparable to other published *de novo* assemblies (Table 1).

To enhance the quality of our genome assembly, we adopted a meta-assembly strategy in which multiple assemblies were merged to yield a single assembly. In meta-assembly strategies, individual assemblies are aligned, and one best assembly is selected for each aligned segment based on the absence of rare SVs, unresolved sequences, or possible mis-assembly inferred by other experimental evidence, such as mate-pair sequencing data^25^. For meta-assembly, Metassembler software^25^ was applied to 37× mate-pair short reads from the three subjects in sum to infer discordance among the individual scaffolds (Supplementary Table 3). A total of 12 meta-assemblies, or sets of meta-scaffolds, were generated from the three sets of scaffolds, based on the order and combination of the processed sets of scaffolds (see Methods section). Among the 12 possible combinations, we found that one combination (jg1c + (jg1a + jg1b))—which merged the scaffolds of jg1c with the meta-scaffolds generated from that of jg1a and jg1b in this order—exhibited no apparent large chimeric mis-assembly in any autosomes. This combination was chosen for the downstream sophistications; the absence of chimeric mis-assembly was assessed using STS markers described later. This set of meta-scaffolds exhibited better contiguity and accuracy than the original set of scaffolds for subject jg1c (Table 1).

Although meta-scaffolds were more contiguous and accurate than individual sets of scaffolds, rare reference alleles should still be retained in the meta-scaffolds. To eliminate these rare reference alleles, we aligned the three individual sets of scaffolds against the meta-scaffolds, performed variant calling, defined the major allele among the three sets of scaffolds, and substituted the minor allele on the meta-scaffolds to the major allele both in terms of SNVs and SVs (Supplementary Fig. 6a). For tri-allelic sites, we chose one allele randomly among the three as a reference allele. We also found that two assemblies among the three contained a 2.6-Mb inversion in the long arm of chromosome 9 (Supplementary Fig. 2), and we confirmed that the meta-scaffolds also contained the inversion.

We next tried to anchor the majority-voted meta-scaffolds on each chromosome. To do so, we utilized a total of 85,386 distinct STS markers from three genetic maps and six RH maps pre-dated the reference genome: the Genethon^26^, deCODE^27^, and Marshfield^28^ genetic maps and the Whitehead-RH^29^, GeneMap99-GB4^30^, GeneMap99-G3^30^, Stanford-G3^31^, NCBI_RH^32^, and TNG^33^ RH maps. We searched for STS marker amplifications by electronic PCR analysis of the meta-scaffolds and used ALLMAPS software^34^ to order and orient the meta-scaffolds to build chromosomes. The co-linearities between the anchored meta-scaffolds and genetic and RH maps were 0.999 ± 0.004 and 0.986 ± 0.021, respectively (Pearson’s correlation coefficient; mean ± SD). However, we found that ALLMAPS using all nine above-mentioned maps did not assign any meta-scaffolds to the Y chromosome, probably because most of the maps did not include the Y chromosome. Nonetheless, we found that ALLMAPS using three of the nine maps (deCODE, TNG, and Stanford-G3) assigned some meta-scaffolds to the Y chromosome as well as autosomes and the X chromosome. Therefore, we adopted the ALLMAPS assignment with the nine maps for autosomal assignment and those with three maps to the sex chromosomes.

After anchoring these meta-scaffolds to chromosomes, we found a chimera in the sex chromosomes. A meta-scaffold harboring the *SRY* locus, a gene on the Y chromosome, was chimeric and anchored to the long arm of the X chromosome in the selected set of meta-scaffolds. We therefore chose a set of meta-scaffolds from another set of meta-scaffolds (jg1a + (jg1b + jg1c)) for the long arm of the X chromosome that had no apparent chimeric region.

We also manually modified the length of unresolved regions in the telomeric, centromeric, and constitutive heterochromatic regions represented as a stretch of Ns (see Methods section). We then masked a pseudo-autosomal region (PAR) in the Y chromosome to guarantee that the resulting sets of sequences represented a haploid. In addition, we shifted the start position of the mitochondrial meta-scaffold to match the revised Cambridge Reference Sequence (rCRS) coordinates^35^, which provides the reference coordinate system for the mitochondrial genome.

The procedure described above yielded a set of chromosome-level sequences for 22 autosomes, 2 sex chromosomes, and 1 mitochondrial chromosome, along with 599 unplaced scaffolds, and we designated this set of sequences JG1 (Figure 1b). The total length of JG1 was approximately 3.1 Gb (Table 1). Notably, in the JG1 genome assembly, 19 chromosomal arms were successfully represented as single scaffolds (Figure 1b). After constructing these chromosome-level sequences, we then aligned them to reference genome GRCh38 using minimap2 software^36^ and found an overall high similarity between the two genomes at the sequence level (Figure 1c). Because JG1 was built independently from the reference genome GRCh38, this overall high similarity provided strong support for our approach for building JG1 described above.

### Representativeness of the JG1 haplotype in terms of SNVs

To assess whether JG1 is a representative reflection of the SNV composition of the Japanese population, we performed PCA using JG1 and the reference hg19, along with 2,022 haplotypes constructed from 11 HapMap3 populations (see Methods section). The PCA plot shows three major clusters representing African, Asian, and European populations (Figure 2a). The JG1 haplotype localized near the cluster of Asian populations, whereas the hg19 haplotype localized between the African and European populations, as expected based on the donors’ ancestries (Figure 2a). Notably, the JG1 haplotype did not localize within the Asian cluster; instead it was the most distant site both from the European and African populations, suggesting an “Asianness” when compared with the other two populations. We also performed PCA with JG1 and 505 haplotypes constructed from three Asian populations: Japanese (JPT), Han Chinese (CHB) and Chinese in Denver (CHD) (Figure 2b). The PCA plot included two distinct clusters (namely, Japanese and others), with the JG1 haplotype associated with the Japanese cluster.

**Figure 2.**
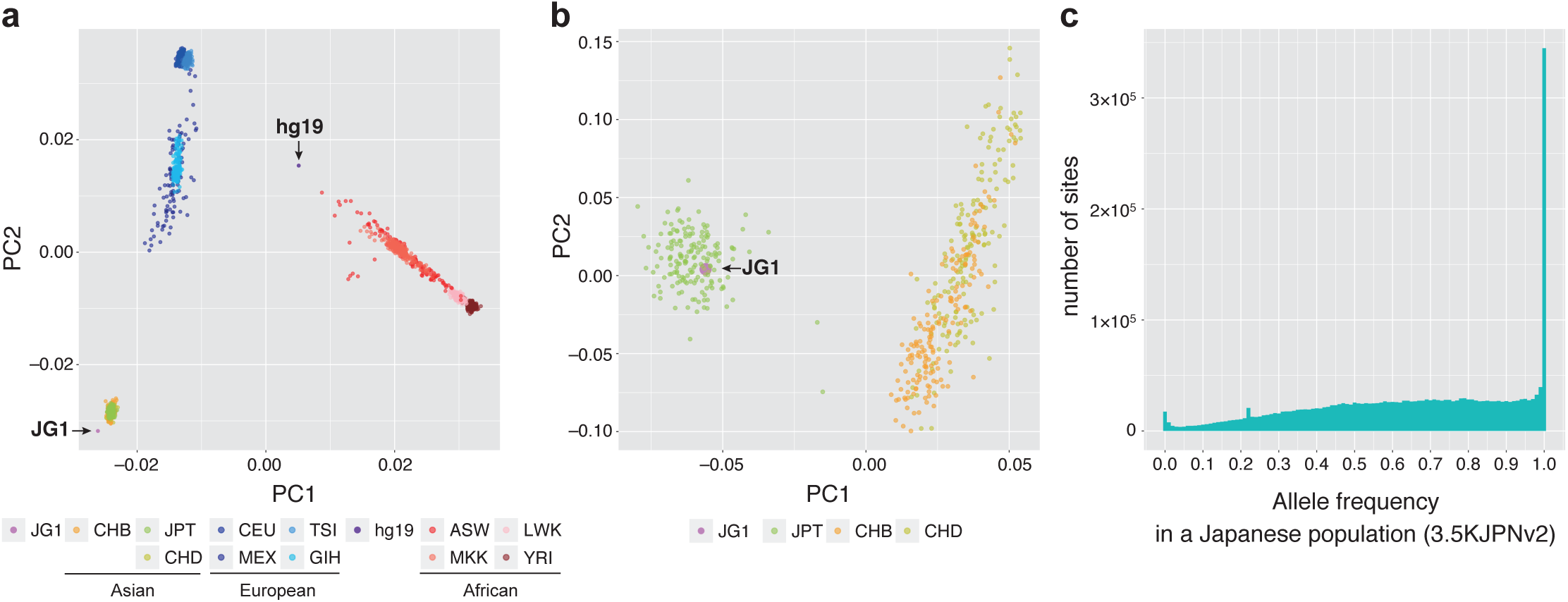
SNV characteristics of JG1. **(a)** PCA plot of the haplotype SNP composition of JG1, the reference hg19, and HapMap3 samples. **(b)** PCA plot of the haplotype SNP composition of JG1 and Asian samples from HapMap3. **(c)** Unfolded site frequency spectrum representing the frequencies of alleles employed in the JG1 sequence in the Japanese population of 3.5KJPNv2.

To assess whether JG1 harbors the major allele among the Japanese population across SNV sites, we first aligned JG1 against the reference genome hs37d5 and detected SNVs. The genome-by-genome alignment and comparison by minimap2 and paftools software^36^ called 2,501,575 SNVs between hs37d5 and JG1 in the autosomes and X chromosome. We then extracted the frequency of the allele employed in JG1 from the allele frequency (AF) panel of 3,552 Japanese individuals (namely, 3.5KJPNv2 AF panel^37^) to create a site frequency spectrum, in which the horizontal axis indicates the non–hg19-type allele and the vertical axis indicates the number of such SNV sites (Figure 2c). From these data, we found 241,500 SNV sites with an AF = 1.0, indicating that all of the Japanese chromosomes in the AF panel carried the JG1-type allele at the 241,500 sites. This corresponds to 97.99% of all such SNV sites that had an AF = 1.0 (246,464) in the 3.5KJPNv2 AF panel. Similarly, we identified 367,271 and 626,254 SNV sites with an AF ≥ 0.99 or ≥ 0.90, respectively, corresponding to 97.11% and 96.24% of such SNV sites in the 3.5KJPNv2 AF panel, respectively (378,211 and 650,718). A peak observed at an AF of ∼0.22 was associated with the SNPs clustered in the XTR region—a region known to harbor complex duplications—within 88.8 to 92.4 Mb on the X chromosome. A peak at an AF of approximately zero could most likely be attributed to artificial SNVs called at the edges of alignments. We also assessed the effectiveness of the majority decision approach. Of the 2,501,575 SNVs, 1,176,922 (47%) and 1,204,762 (48%) were detected in three and two of the three JG1-donor individuals, respectively (Supplementary Fig. 6b).

### Representativeness of the JG1 haplotype in terms of SVs

To investigate differences between JG1 and GRCh38 in terms of SVs, we aligned JG1 against the reference GRCh38 and detected SVs (insertions and deletions) using the minimap2 and paftools software programs^36^. A genome-by-genome comparison detected 8,689 insertions and 6,177 deletions >50 bp but <10,000 bp in length. The length distribution of the SVs exhibited two peaks, at approximately 300 bp and 6 kb (Figure 3a). We confirmed that the 300-bp and 6-kb peaks were associated with *Alu* and LINE1, respectively. Most of the SV-associated *Alu* and LINE1 were classified as *Alu*Y and L1HS, respectively, both of which constitute the currently-active subclass of these transposable elements (the length distributions of detected transposable elements are shown in Supplementary Fig. 7). In addition, the detected SVs were often observed in the sub-telomere–telomere regions (Figure 3b), consistent with a previous report^12^.

**Figure 3.**
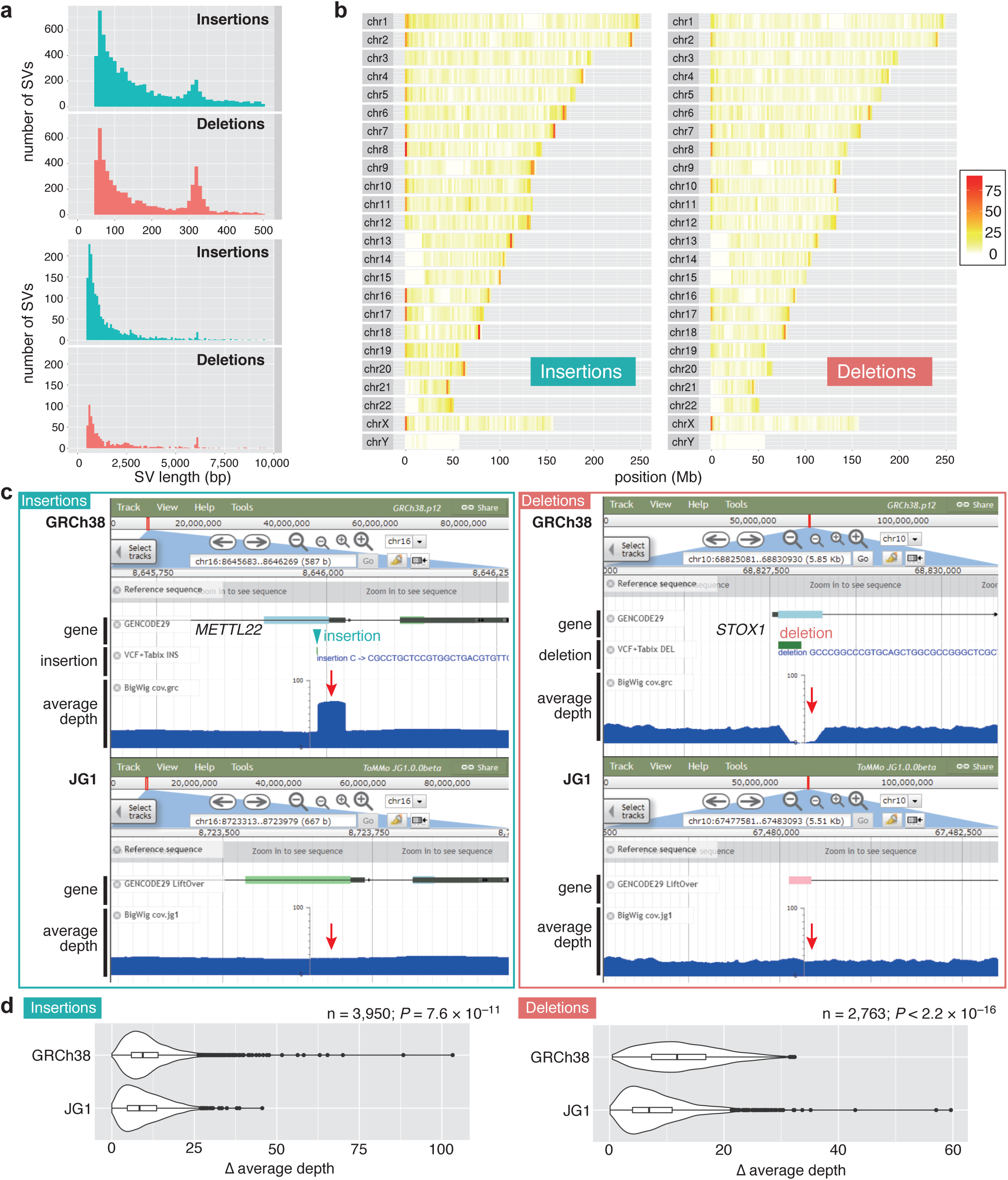
Analysis of JG1 SV. **(a)** Length histogram of small (≤500 bp) and large (>500 bp) insertions and deletions detected by comparing JG1 and GRCh38. **(b)** Distribution of insertions and deletions among the chromosomes of GRCh38. **(c)** JBrowse snapshots of one insertion/deletion example. Tracks are GENCODE gene annotations, detected SVs, and average depth of short reads from 200 Japanese samples. **(d)** Difference in average depth between the SV and upstream regions of same length as the SV for GRCh38 and JG1.

To investigate the extent to which JG1 represents a Japanese population in terms of SVs, we mapped short reads of 200 Japanese individuals to JG1 and GRCh38 to compare the average read depth among the 200 individuals around the SVs in JG1 and GRCh38. As shown in Figure 3c, insertions were typically associated with a ‘piling-up’ of the average depth, whereas deletions were typically associated with a depression of the average depth in GRCh38. Neither pattern was clearly evident in the corresponding region in JG1 (Figure 3c), suggesting that most of the Japanese samples shared the SVs. To determine whether this pattern is common among SVs throughout the genome, we compared the maximum difference in average depth between the SV region and its adjacent upstream region of the same length and found that the difference in the average depth was smaller in JG1 than GRCh38 (Figure 3d; *P* = 7.6 × 10^-11^ for n = 3,950 pair of insertions; *P* < 2.2 × 10^-16^ for n = 2,763 pair of deletions; Wilcoxon signed rank tests).

### Utility of JG1 as a reference for rare disease exome analyses

To evaluate whether JG1 is a suitable reference genome for clinical NGS analyses, we examined exomes of Japanese families with rare diseases^38^. The sample cohort consisted of 22 individuals from six trio families and one quartet family. All of the families had one child affected with diplegia and eight causal variants were identified in previous analyses using the reference genome hg19 (Table 2). The diseases exhibited autosomal recessive, compound heterozygous, and autosomal dominant modes of inheritance, including *de novo* mutations, and the causal variants included both single nucleotide and deletion variants.

**Table 2.** Diplegia cohort.

| **ID\*** | **family** | **type** | **locus** | **variant(s)** | **variant effect prediction** | **mode of inheritance** |
| --- | --- | --- | --- | --- | --- | --- |
| 3 | FD-05 | trio | *CTNNB1* | c.1683+2T>C | HIGH (splice donor) | *de novo* SNV |
| 5 | FD-07 | trio | *CYP2U1* | c.651delC | HIGH (frame shift) | autosomal recessive deletion |
| 6 | FD-08 | trio | *SPAST* | c.1276C>T | MODERATE (missense) | *de novo* SNV |
| 7 | FD-09 | quartet | *GNAO1* | c.736G>A | MODERATE (missense) | *de novo* SNV |
| 9 | FD-11 | trio | *CACNA1A* | c.653C>T | MODERATE (missense) | *de novo* SNV |
| 10 | FD-12 | trio | *SPAST* | c.1496G>A | MODERATE (missense) | *de novo* SNV |
| 11 | FD-13 | trio | *AMPD2* | c.515+1G>A<br>c.1724C>T | HIGH (splice donor)<br>MODERATE (missense) | compound heterozygous SNV |

\* Case ID from Table 2 in Takezawa et al<sup>38</sup>.

To facilitate exome re-analysis with JG1, we lifted over the resource bundles of Genome Analysis ToolKit (GATK) software (which are used for accurate variant calling) and GENCODE gene annotation information^39^ to predict variant effects (see Methods section) and performed exome analyses according to GATK best practices. The JG1-based exome analyses correctly identified all (8/8) of the previously reported causal variants. In addition, the total number of called variants was lower in JG1 than the reference hs37d5 (Figure 4a). This comparison was done in the 225,888 exome regions with one-to-one correspondence between JG1 and hs37d5 (87,971,409 bp and 87,997,786 bp for JG1 and hs37d5, respectively). Moreover, the number of both high- and moderate-impact variants (which are the primary causal variant candidates) was also lower in JG1 than hs37d5 (Figure 4b; 473 ± 16 vs 671 ± 13 high-impact and 8,774 ± 97 vs 10,599 ± 89 moderate-impact variants for JG1 and hs37d5, respectively; mean ± SD). These findings suggest that JG1 produces fewer false-positives while successfully detecting disease-causing variants in whole-exome analyses. In addition, we compared the variants detected with JG1 and hs37d5 by lifting over the JG1-detected variants to hs37d5 and found ∼15,000, ∼29,000, and ∼52,000 specific to JG1, hs37d5, and both references, respectively (Figure 4c). Moreover, we extracted the non–GRC-type AF in the JG1-specific, hs37d5-specific, and shared variant sites from the 3.5KJPNv2 AF panel and found that most of the hs37d5-specific variants were major alleles among the Japanese population, whereas the shared and JG1-specific variants tended to be biased toward the minor AFs (Figure 4d).

**Figure 4.**
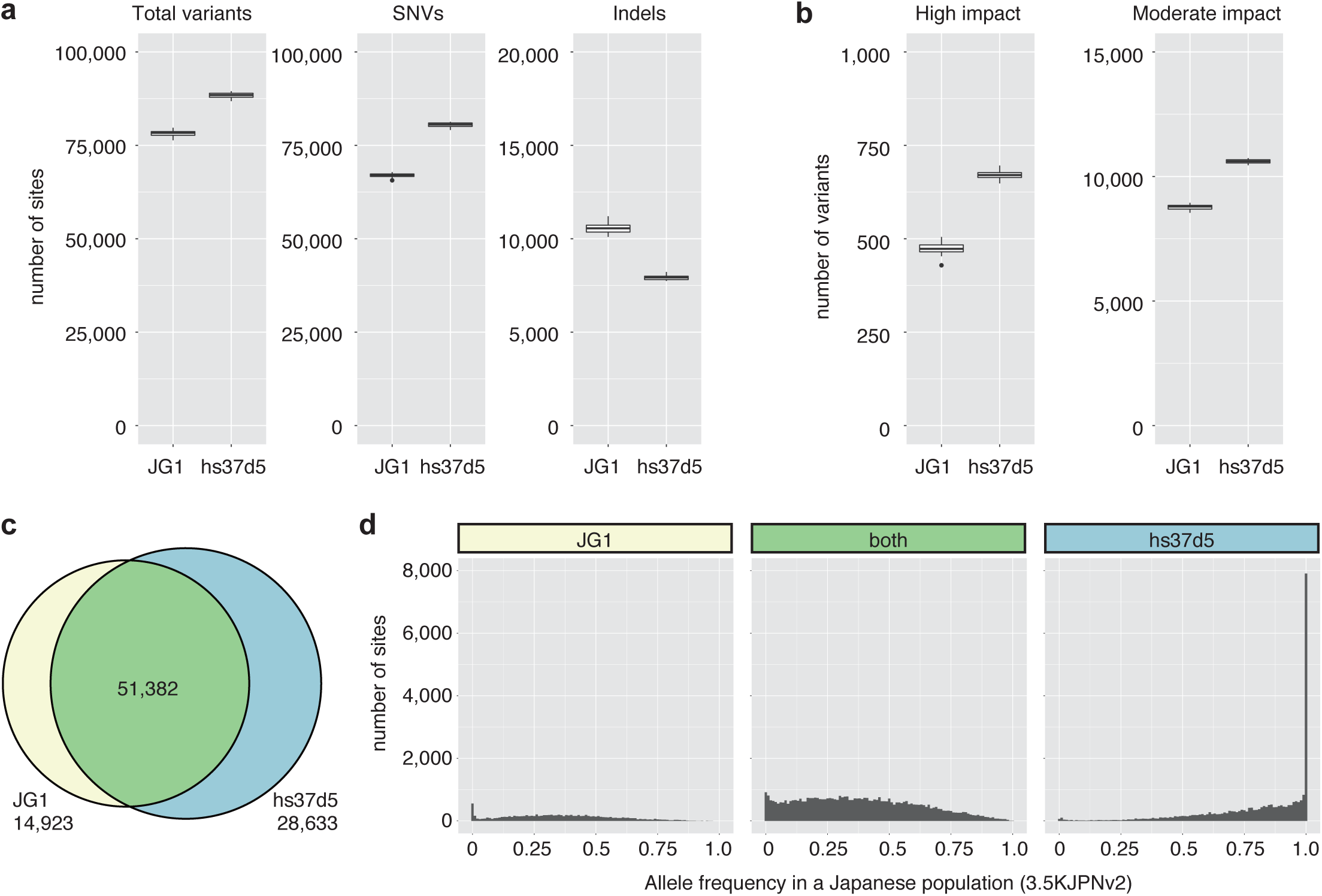
Comparison of variants called in exome analyses employing JG1 or hs37d5 as a reference genome. (**a**) Number of total variants, SNVs, and short indels called per individual. (**b**) Number of high- and moderate-impact variants. **(c)** Venn diagram showing overlap relationships between variants detected in JG1 (lifted over to the hs37d5 coordinates) and those detected in hs37d5. Shown are results for a representative individual. **(d)** Unfolded site frequency spectra representing the frequency of non–GRC-type alleles in the variant sites detected specifically in JG1, in both genomes, and specifically in hs37d5, respectively. Shown are results for the same individual as in **(c)**.

## DISCUSSION

Here, we report the first construction of a Japanese haploid genome sequence, JG1, by integrating three highly contiguous *de novo* hybrid assemblies from three Japanese donor individuals to build a population-specific (i.e. ethnicity-matched) reference genome. Employing a meta-assembly approach produced a more-contiguous and accurate assembly, and relying on majority decision among the three genomes substituted most of the rare reference alleles. The results of both SNV and SV analyses suggested that the JG1 haplotype represents major variation among the Japanese population. Moreover, we demonstrated that JG1 exhibits several advantages as an ethnicity-matched reference for NGS analyses, at least within the clinical context of whole exomes of Japanese samples. Using JG1 could thus facilitate detecting the proverbial needle in a haystack, by reducing the size of the haystack in NGS analyses of the Japanese population.

Integration and majority decision regarding multiple assemblies to yield a single haploid genome can produce a highly contiguous and accurate assembly, thus effectively eliminating most rare reference variations. Haploid representation of the genome is beneficial because it is compatible with many conventional bioinformatics tools developed to date for mapping, variant calling, predicting variant effects, and subsequent interpretations. Although we appreciate that the development of a pan-human genome graph could be the next milestone reached in comprehensively assessing human genetic variations among diverse populations, we expect that population-specific reference genome such as JG1 will prove to be practical and beneficial options for genome analyses of individuals originated from the population.

Several limitations of the current version of JG1 should be noted: (1) sequence incompleteness and gaps/un-localized fragments remaining, which could result in erroneous mapping and variant calling; (2) few original annotations on the JG1 coordinates; and (3) incomplete representation of the major variations in the Japanese population. The incompleteness of the genome sequence could be largely overcome by applying other genome sequencing technologies, including ultra-long reads of Oxford nanopore technologies in combination with targeted cloning from whole-genome BAC libraries. Chromosome-scale scaffolding using Hi-C^40^ could also contribute to the generation of more-contiguous assemblies. The genome of a Japanese complete hydatidiform mole, characterized by a duplicated haploid genome, could also contribute to gap-filling due to ease of assembly^41^. The limitation of few original annotations could be overcome by constructing an AF panel with JG1 as the reference and by *de novo* prediction or experimental inference regarding gene regions. More comprehensive lifting-over of many annotations would also be practically important. The representativeness of the major alleles would be improved by adding more assemblies. One approach that could be used for addition is the phased diploid assembly^23^, which provides a pair of haplotype (i.e., diplotype) assemblies from a single individual. Because the two haploid genomes can be regarded as a random sample from a panmictic population, assembling two haploid genomes per individual can increase the representativeness of variations in the population. Despite its limitations above, the current version of JG1 represents a useful tool for efficient causal variant detection.

Additional samples, for example hundreds of samples from a single population, would be beneficial for constructing population-specific reference genomes in the future, not only with respect to SNVs but also SVs, although less is known regarding the entire repertoire of SVs present in a population than that of SNVs. Both integrative haploid reference genomes such as JG1 and collective genome reference developed in the future such as genome graphs—both of which can be constructed from hundreds of *de novo* assemblies—should advance the accuracy of genome analyses and facilitate development of personalized medicine approaches.

## METHODS

### Selection and analysis of donor individuals

#### Donor selection

Three male Japanese volunteers were recruited and participated in this study with written, informed consent.

#### G-banding analysis (Supplementary Fig. 1a–c)

G-banding analyses for the three volunteers were performed using phytohemagglutinin-stimulated lymphocytes at the laboratory of SRL Inc. (Tokyo, Japan).

#### PCA of donors with Japanese samples (Fig. 1a)

Paired-end reads with length of 162 bp from the three donors (jg1a, jg1b, and jg1c) were individually mapped to hs37d5.fa, and variant calling was performed according to previously described methods^37^, following GATK Best practices. The resulting VCF file was subjected to PCA using EIGENSOFT software (ver. 4.2). We chose 310 Japanese samples from the 3.5KJPNv2 cohort^37^; 100 from Miyagi Prefecture in northern Japan; 29 from Nagahama City, in western Japan; and 181 from Nagasaki Prefecture, in southern Japan. Variants shared among the 313 samples were selected and filtered using plink software (ver. 1.9) with the ‘--geno 0.05 --maf 0.05 --hwe 0.05’, and ‘--indep-pairwise 1500 150 0.03’ options. The resulting dataset consisted of 18,658 variants.

### Genome analyses

#### Long-read SMRT sequencing

Long-read SMRT sequencing was performed as previously described^19^. Briefly, genomic DNA from nucleated blood cells was sheared to ∼20 kb and used for library preparation with a DNA template prep kit 2.0 (Pacific Biosciences; Menlo Park, CA). Size selection was carried out using the Blue Pippin system (Sage Science; Beverly, MA), targeting 18 kb (10-15 kb for some libraries of jg1a). The libraries were sequenced on a PacBio RSII instrument using P6-C4 chemistry.

#### Optical mapping

Optical mapping was performed using Irys system or Saphyr system, according to the manufacturer’s protocol (Bionano Genomics; San Diego, CA). For sample jg1a, high-molecular-weight genomic DNA from nucleated blood cells was nicked using the endonucleases Nt.BspQI or Nb.BssSI and then labeled with fluorophore-tagged nucleotides. The labeled DNA was imaged on the Irys system. For samples jg1b and jg1c, high-molecular-weight genomic DNA from nucleated blood cells was labeled using direct labeling and staining (DLS) chemistry. The labeled DNA was imaged on the Saphyr system.

#### Short-read paired-end sequencing

Short-read paired-end sequencing was performed as previously described^42^. Briefly, genomic DNA from buffy coat samples was fragmented to an average target size of 550 bp, and then subjected to library construction using a TruSeq DNA PCR-Free HT sample prep kit (Illumina; San Diego, CA). The libraries were sequenced on a HiSeq 2500 system (Illumina) with a TruSeq Rapid PE Cluster kit (Illumina) and TruSeq Rapid SBS kit (Illumina) to obtain 162- or 259-bp paired-end reads.

#### Mate-pair sequencing

Genomic DNA from nucleated blood cells was used for library construction with a Nextera Mate Pair Library Preparation kit, gel-free protocol (Illumina). The libraries were size-selected to an average of 500 bp using AMPure XP beads (Beckman Coulter; Indianapolis, IN) and sequenced on a HiSeq 2500 system (Illumina) with a TruSeq Rapid PE Cluster kit (Illumina), and TruSeq Rapid SBS kit (Illumina) to obtain 201-bp paired-end reads.

### Overview of the computational methods for JG1 construction

A diagram showing an overview of the construction of JG1 is provided in Supplementary Fig. 3. JG1 was constructed according to the following steps, which are also described in the download page for the JG1 sequence file from the jMorp website (https://jmorp.megabank.tohoku.ac.jp/dj1/datasets/tommo-jg1.0.0.beta-20190424/files/tech-notes-for-computation.pdf). The computation was carried out by using the Tohoku Medical Megabank Organization (ToMMo) Super Computer (https://sc.megabank.tohoku.ac.jp/en/outline).

#### *De novo* assembly of PacBio subreads

PacBio subreads were assembled using Falcon software^23^ (build ver. falcon-2017.11.02-16.04-py2.7-ucs2.tar.gz) with the following configurations: reads shorter than 9 kb were used for error-correction of the longer reads (’length_cutoff = 9000’), and error-corrected reads longer than 15 kb were used for assembly (’length_cutoff_pr = 15000’). Detailed settings are provided below:

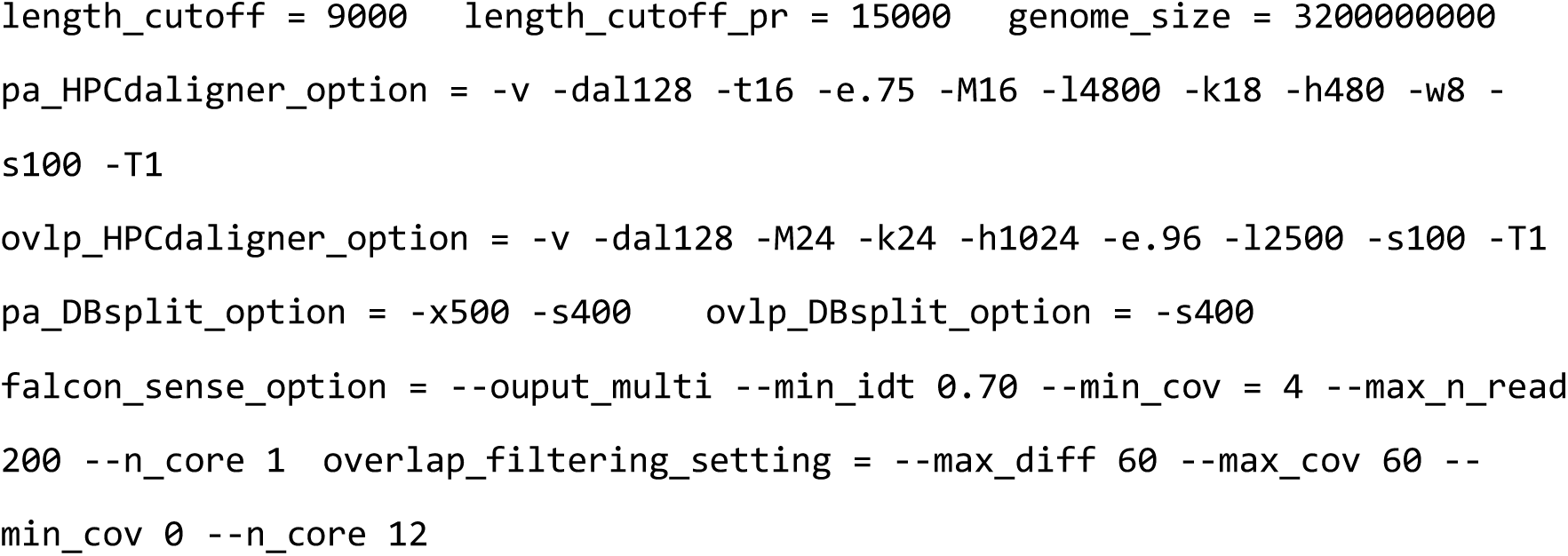

The contigs were then polished with the PacBio subreads using ArrowGrid software^24^ (ver. 81b03f1; GitHub commit tag), with slight modifications to accommodate our number of data files and UGE settings.

#### *De novo* assembly of Bionano optical maps

We obtained two sets of Bionano data using two different enzymes, Nt.BspQI and Nb.BssSI, for subject jg1a, and one set of Bionano data was obtained with DLE-1 for jg1b and jg1c. In both cases, the Bionano data were assembled in two steps—a rough assembly step and a full assembly step—to perform *de novo* assembly as independently as possible from the reference. For the rough assembly step for jg1a, we ran pipelineCL.py software using the following settings:

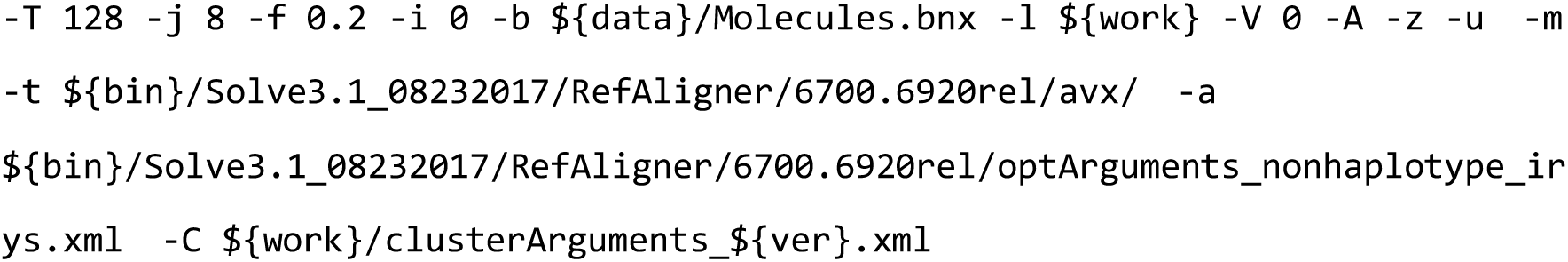

For the full assembly step for subject jg1a, we ran the software using the following settings:

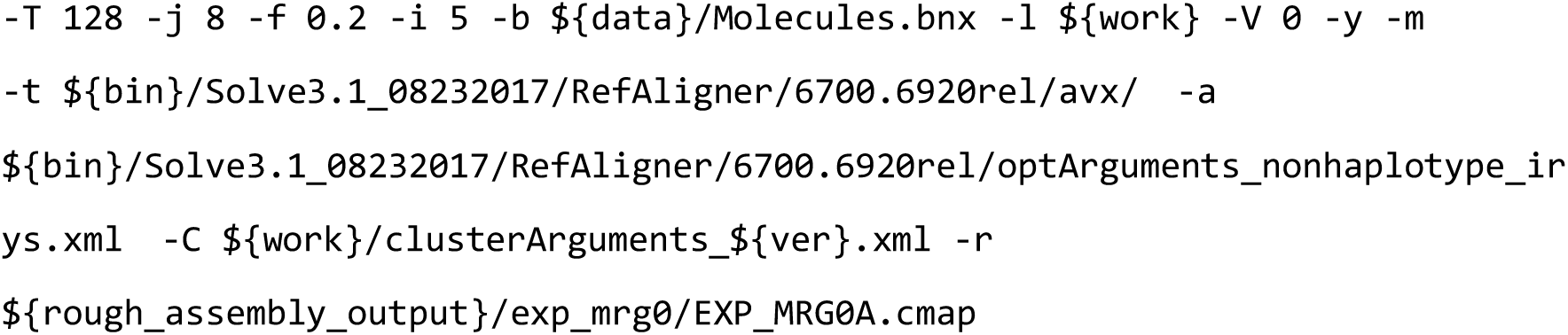

For the rough assembly step for subjects jg1b and jg1c, we ran the software using the following settings:

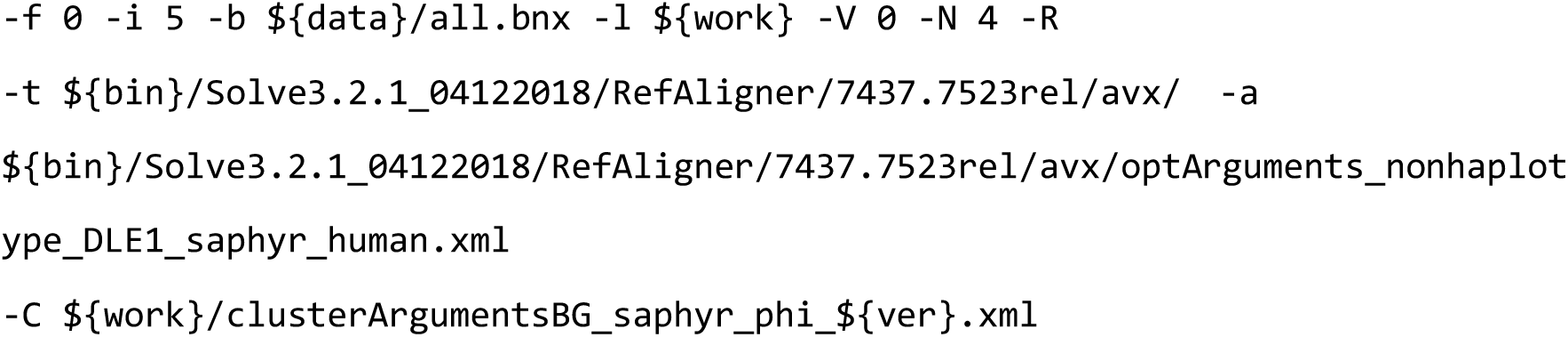

For the full assembly step of subjects jg1b and jg1c, we ran the software using the following settings:

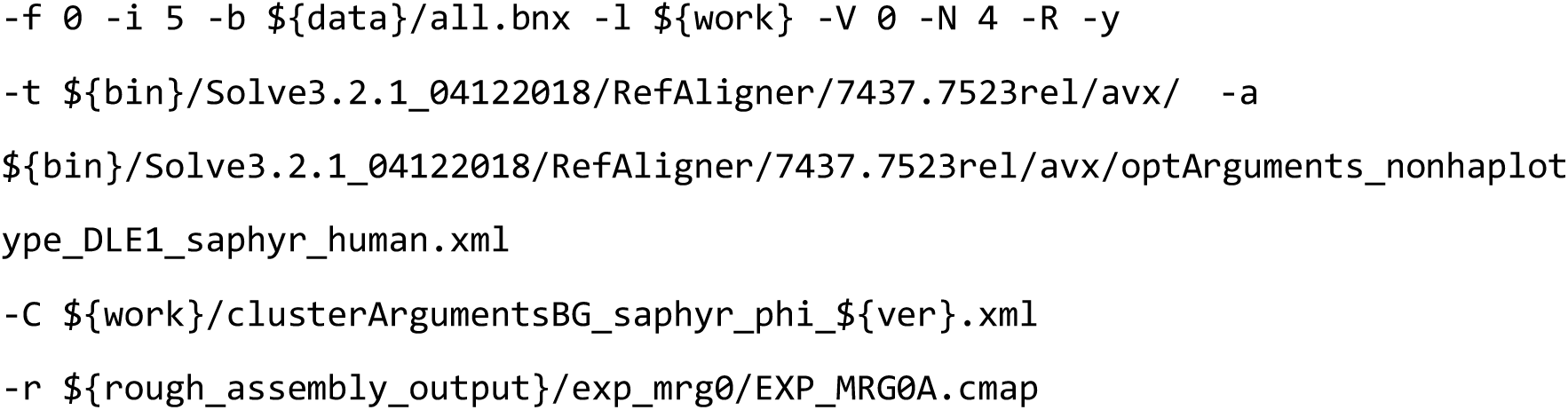

The ‘-T’ and ‘-j’ options were varied for computational efficiency. The BionanoSolve software suite was used for the above computation. We used BionanoSolve (ver. 3.1) for the assembly for subject jg1a, and ver.3.2 for the assembly for subjects jg1b and jg1c.

#### Hybrid scaffolding

Hybrid scaffolding was performed using BionanoSolve software (ver. 3.2). Hybrid scaffolding for subject jg1a was performed in the two-enzyme hybrid scaffolding mode using the runTGH.R script with the following options:

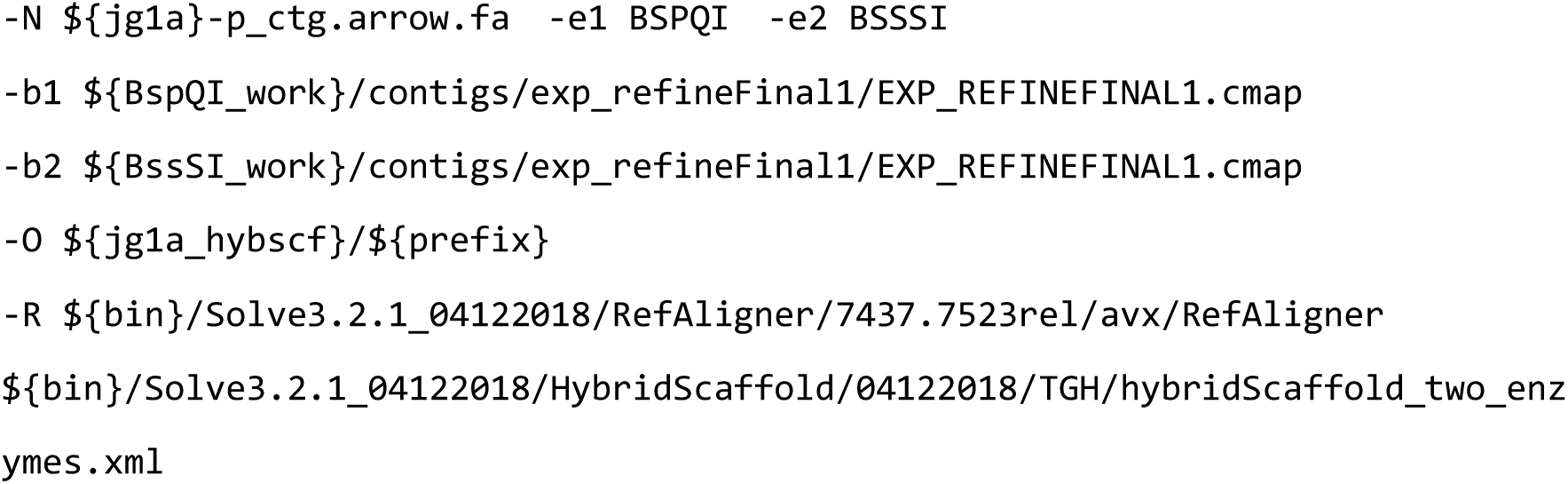

Hybrid scaffolding for subjects jg1b and jg1c was performed in the single-enzyme hybrid scaffolding mode, using the hybridScaffold.pl script with the following options:

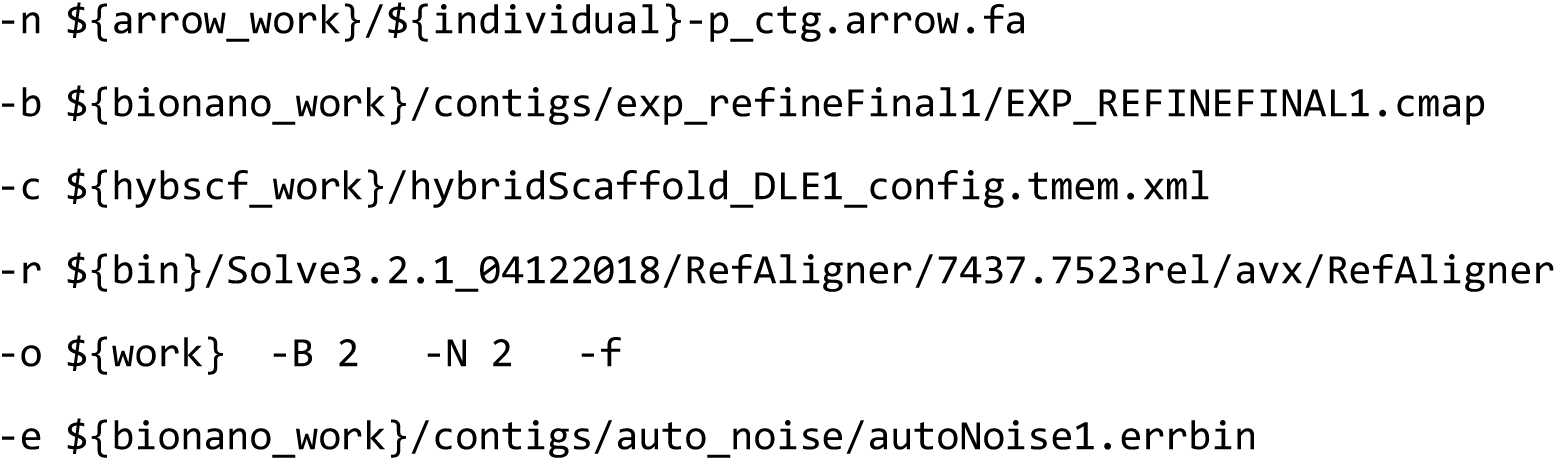

#### Error correction with short reads

Two sets of Illumina paired-end short reads with lengths of 162 bp and 259 bp were mapped to the hybrid scaffolds using BWA-MEM software^4^ (ver. 0.7.17) with the option ‘-t 22 -K 1000000’. The alignment file was coordinate-sorted and compressed using the Picard tools (ver. 2.18.4) SortSam command. The resulting BAM files for the 162- and 259-bp paired-end reads were merged using the Picard tools MergeSamFiles command. The merged BAM files were then split to each scaffold using SAMtools^43^ (ver. 1.8) view command, and then each scaffold was polished using Pilon software^44^ (ver. 1.22, modified to correct the issue reported at https://github.com/broadinstitute/pilon/issues/48) with the option ‘--threads 22 --diploid --changes --vcf --tracks’. The FASTA files for each polished scaffold were then merged into a single multi-FASTA format file.

#### Meta-assembly

The three sets of polished scaffolds were then meta-assembled using Metassembler software^25^ (ver. 1.5; with modification of the type of ‘totalBases’ variable in the CEstat.hh from int to long to accommodate large genomes). There were 12 possible combinations to meta-assemble the three sets: (a + (b + c)), (a + (c + b)), ((a + b) + c), ((a + c) + b), (b + (a + c)), (b + (c + a)), ((b + a) + c), ((b + c) + a), (c + (a + b)), (c + (b + a)), ((c + a) + b), and ((c + b) + a), where x + y indicates meta-assemble x and y in this order. For each round of meta-assembly, we aligned the two assemblies using the NUCmer command of MUMmer software^45^ (ver. 4.0.0beta2) with the option ‘--maxmatch -c 50 -l 300’. The resulting DELTA file was filtered using delta-filter software with the option ‘-1’ to extract one-to-one correspondence. Next, the DELTA file was converted to COORDS format using the show-coords command with ‘-clrTH’ option. Short mate-pair reads were classified into four categories (mp, pe, se, and unknown) using NxTrim software^46^ (ver. 0.4.3), and the resulting set of reads with the correct mate-pair orientation (mp) were mapped using Bowtie2 software^47^ (ver. 2.3.4.1) with the ‘--minins 1000 -- maxins 16000 --rf’ options. The output SAM file was then processed using the mateAn command with ‘-A 2000 -B 15000’ option, indicating that the range of insert length was 2 to 15 kb. The NUCmer alignment information and the mate-pair mapping information were integrated using the asseMerge command with ‘-i 5 -c 6’ option. Finally, the resulting METASSEM file was converted to FASTA format using the meta2fasta command.

#### Major allele substitution

The three sets of polished hybrid scaffolds were aligned to the 12 sets of meta-scaffolds using minimap2 (ver. 2.12), and variants were called using the paftools call command. After normalizing the manner of variant representation using the BCFtools norm command (ver. 1.8), SNVs and SVs shared by two of the three genomes were detected using the BCFtools isec command, and these were regarded as the major alleles and employed in JG1 using the BCFtools consensus command. For multi-allelic sites, one allele was chosen randomly.

#### Detection of STS marker amplification by electronic PCR

We detected amplification of the STS markers in the three genetic and six RH maps (Genethon, Marshfield, and deCODE genetic maps; GeneMap-G3, GeneMap99-GB4, TNG, NCBI_RH, Stanford-G3, and Whitehead-RH maps) from the meta-scaffolds using the in-house electronic PCR software gPCR (ver. 2.6a) with the ‘-S -D’ option (’-S’ to show amplicon sequence, ‘-D’: to show direction of markers). The STS markers were obtained from the UniSTS database (ftp://ftp.ncbi.nih.gov/pub/ProbeDB/legacy_unists/). The results were used to infer the presence of chimeric scaffolds. One set of meta-scaffolds (jg1c + (jg1a + jg1b)) was selected for the primary downstream analysis. In addition, to build the X and Y chromosomes, an additional set of meta-scaffolds (jg1a + (jg1b + jg1c)) was selected.

#### Anchoring scaffolds to chromosomes

The electronic PCR results were converted to BED format files, and the coordinates of some RH maps were scaled to approximately 2,000 to fit those for the genetic maps; this was done to make it easier to understand the visualization results of the ALLMAPS software^34^ (ver. 0.8.12) but did not affect the anchoring results. These maps were merged using the ALLMAPS mergebed command, and then processed using the ALLMAPS path command with the option ‘-- gapsize=10000’ to anchor the meta-scaffolds to the chromosomes. The weights of each of the three genetic maps was set to 5, and that of each of the RH maps was set to 1 in the weights.txt file. To anchor the sex chromosomes, three maps (deCODE, TNG, and Stanford-G3) that could anchor some scaffolds to the Y chromosome were used.

#### Manual modification

The physical lengths of the short arms of acrocentric chromosomes 13, 14, 15, 21, and 22 were obtained from Table 4 of Morton (1991)^48^. The relative length estimates of constitutive heterochromatin regions in the chromosome 1, 9, 16 were obtained from ref. 49 and ref. 50. The relative length estimate of heterochromatin segment of the Y chromosome was obtained from ref. 51–53. These relative lengths were converted to the base-pair length (Mb) by using the chromosomal arms shown in Table 4 of Morton (1991)^48^. The length of consecutive Ns for each chromosome is provided in Supplementary Table 5. For all chromosomes except 8 and 11, 3-Mb consecutive Ns were inserted instead of 10-kb Ns inserted by the ALLMAPS software, between the two scaffolds flanking the centromere. For chromosomes 8 and 11, in which the centromere-specific sequence repeats were identified in the midst of a scaffold by aligning the LinearCen1.1 sequences^54^ using minimap2 software, no centromeric Ns were inserted. The position of the centromere was inferred from the Whitehead-RH and GeneMap99- GB4 maps, in which the centromeric or constitutive heterochromatin region could be inferred from the region sparsely covered by STS markers, possibly due to the radiation conditions.

#### Building the X and Y chromosomes

We noted that one set of meta-scaffolds, (jg1c + (jg1a + jg1b)), contained a chimeric scaffold between the long arm of the X chromosome and the *SRY* locus of the Y chromosome. To reduce the chimeric meta-scaffolds, we chose apparently non-chimeric scaffolds anchored to the long arm of the X chromosome from another set of meta-scaffolds, (jg1a + (jg1b + jg1c)), and linked them to the scaffold of the short arm of the X chromosome.

#### Masking the pseudo-autosomal region

To locate the pseudo-autosomal regions, we aligned both the X and Y chromosomes from JG1 using minimap2 with the option ‘-cx asm5’, and vice versa. The alignment started from the terminal region of the Y chromosome and ended at 2.26 Mb. This region was regarded as the putative PAR1 region. Other regions such as PAR2 and XTR were probably unresolved for unknown reasons. The putative PAR1 region was masked using the BEDTools software^55^ (ver. 2.27.1) maskfasta command.

#### Mitochondrial chromosome

We aligned the set of meta-scaffolds to GRCh38, the mitochondrial sequence of which was obtained from the rCRS using minimap2 with the option ‘-cx asm5’ to identify a scaffold that corresponds to the mitochondrial genome. We found a scaffold of 16,568 bp in length corresponding to the mitochondrial sequence in another set of meta-scaffolds (jg1a + (jg1b + jg1c)). The start site of the scaffold and the rCRS sequence differed; therefore, we shifted the start site of the scaffold to match that of the rCRS sequence.

### Idiogram drawing (Fig. 1b)

Idiograms were depicted using JG1 BED files scaled to 90% of the original length so that the drawing of JG1 chromosomes longer than that of GRCh38 would be successful using the NCBI Genome Decoration Page (https://www.ncbi.nlm.nih.gov/genome/tools/gdp). The length of the chromosomal arms and the centromeric regions of the idiograms were manually modified to fit the scaffold length of JG1.

### Possible shared large inversion (Supplementary Fig. 2)

Two large scaffolds corresponding to chromosome 9 were extracted from each assembly using the faSomeRecords command. Orientation was carried out by using the seqtk software (ver. 1.3) ‘seq -r’ command. Next, the chromosome 9 sequence from GRCh38 and the two large scaffolds from each subject were aligned using minimap2 (ver. 2.12) with the ‘-t 12 -x asm5 --cs’ option. Harr plots were drawn using the minidot command (bundled with miniasm software^56^ [ver. 0.2]) with the ‘-L’ option. The idiogram of chromosome 9 was drawn using the NCBI Genome Decoration Page.

### PCA of the JG1 and hg19 haplotypes with HapMap3 haplotypes (Fig. 2a, b)

Two haplotypes were constructed for each individual of the HapMap3 variant information. JG1 haplotypes were constructed by aligning JG1 to hs37d5 using minimap2 software and identifying the allele at the marker sites. Variants shared among the 2,022 HapMap3 haplotypes and JG1 and hg19 haplotypes were filtered the using plink software with ‘-- geno 0.05 --maf 0.05’ option. The resulting dataset consisted of 178,047 variants. PCA was performed using EIGENSOFT software. For PCA of JG1 and three Asian populations, 505 haplotypes from the JPT, CHB, and CHD populations were chosen. Four CHD samples were omitted due to apparent inconsistency inferred from a pre-analysis of the PCA plots including these samples.

### SV analysis (related to Fig. 3)

#### SV detection

The GRCh38 sequence was downloaded from illumina iGenome website (ftp://ussd-ftp.illumina.com/Homo_sapiens/NCBI/GRCh38/Homo_sapiens_NCBI_GRCh38.tar.gz). The JG1 sequence was aligned to GRCh38 using minimap2 (ver. 2.12) with the ‘-t 24 -cx asm5 --cs=long’ option. The resulting PAF file was subjected to variant calling using the paftools call command. The VCF file was normalized using the BCFtools (ver. 1.8) norm command with the ‘--threads 4 --remove-duplicates’ option. SVs ≥51 bp and <10 kb were subjected to the downstream analyses.

#### Average depth analysis

Two hundred Japanese individuals (100 males and 100 females) were selected from the 3,552 samples^37^. The 162-bp paired-end reads were mapped using BWA-MEM software as described^37^. Next, accessible regions were defined as the regions where the average depth among the 200 individuals is ≥5 and ≤mean + 2SD; mean and SD were computed for each chromosome. SVs detected within the accessible regions and detected by comparing same autosomes of GRCh38 and JG1 were considered. The mean value of the average depth within the adjacent upstream region of SV was regarded as the reference value, and the Δ average depth was defined as the difference between the reference value and the value of the position showing the maximum absolute difference within the SV region.

### Rare disease exome analysis (related to Fig. 4)

Exome analyses were carried out following the GATK best practices for germline variant detection. Short reads were mapped using BWA-MEM software, and the resulting alignment files were sorted and duplication-marked using SAMtools^43^ software. Variants of the disease cohort families were called using GATK software (ver. 4.0 to 4.1), and the joint calling process was carried out with samples from other Japanese subjects with various rare diseases. The BED files describing the exome capture regions (SureSelect Human All Exon V5, Agilent) were lifted over using the paftools liftover command. The GATK resource bundles were lifted over to the JG1 coordinates using the Picard tools LiftoverVcf command. GENCODE (ver. 29) annotations were lifted over to the JG1 coordinates using an in-house script. The chain files, which were required for lifting over, were generated from the results of minimap2 with an in-house script. The SnpEff database^58^ was constructed using the lifted-over GENCODE annotation file and used for variant effect predictions. Variants called against JG1 were lifted over to the hs37d5 coordinates by using the Picard tools LiftoverVcf command. Overlap relationships between the variants were assessed using the BCFtools (ver. 1.9) isec command.

### Statistical tests and graph drawing

Statistical tests were performed using R software (ver. 3.5.1). Histograms were drawn using R software (ver. 3.5.1) and ggplot2 library (ver. 3.0.0).

### Data availability

JG1 sequence, chain files and GENCODE annotation files are available from the jMorp website (https://jmorp.megabank.tohoku.ac.jp/201911/downloads#sequence).

## ACKNOWLEDGEMENTS

This work was supported in part by the Tohoku Medical Megabank (TMM) Project from the Ministry of Education, Culture, Sports, Science and Technology (MEXT) and the Reconstruction Agency; the Japan Agency for Medical Research and Development (AMED; Grant Numbers JP19km0105001 and JP19km0105002) for Tohoku University. All computational resources were provided by the ToMMo supercomputer system (http://sc.megabank.tohoku.ac.jp/en), which is supported by Facilitation of R&D Platform for AMED Genome Medicine Support conducted by AMED (Grant Number JP19km0405001). We appreciate all the volunteers who participated in the TMM project.

## AUTHOR CONTRIBUTIONS

J.T., S.T, K.Y., C.G., T.F., S.M., and Y.O. performed computational analyses. J.T., A.K., S.K., and G.T. interpreted the results of rare disease re-analyses. F.K., J.K., A.O., and J.Y. designed and conducted experiments. J.T. and G.T. wrote the manuscript with the assistance of the others. K.K., M.Y., and G.T. conceived and supervised the project. All authors read and approved the final manuscript.

## SUPPLEMENTARY FIGURE LEGENDS

**Supplementary Fig. 1.** Karyotypes of the three subjects: **(a)** jg1a, **(b)** jg1b, and **(c)** jg1c. The arrow in panel **(a)** indicates the normal variation inv(9)(p12q13).

**Supplementary Fig. 2.** Harr plot of the alignment between chromosome 9 of GRCh38 and two largest scaffolds aligned to chromosome 9 from the **(a)** jg1a, **(b)** jg1b, and **(c)** jg1c assemblies, indicating that the two individual genomes harbor a possible shared inversion. ‘Super-scaffold’ is the default prefix designated by BionanoSolve software.

**Supplementary Fig. 3.** Workflow of the construction of JG1. **(a)** Workflow of the construction of each draft assembly. **(b)** Workflow of the integration of three draft assemblies. Rectangles indicate substrates such as reads, contigs, and scaffolds. Rectangles with rounded corners indicate software or processes.

**Supplementary Fig. 4:** Histogram of PacBio subreads length for **(a)** jg1a, **(b)** jg1b, and **(c)** jg1c. The length of each subread was calculated using the SAMtools (ver. 1.8) faidx command.

**Supplementary Fig. 5:** Histogram of Bionano data for **(a)** Nt.BspQI of jg1a, **(b)** Nb.BssSI of jg1a, **(c)** jg1b, and **(d)** jg1c. The length of each molecule was extracted from the BNX file.

**Supplementary Fig. 6:** Majority decision. **(a)** Schematic representation of majority decision approach. **(b)** Venn diagram of SNVs detected in JG1, jg1a, jg1b, and jg1c by comparison with hs37d5. The intersection relationship was inferred using the BCFtools (ver. 1.8) isec command.

**Supplementary Fig. 7:** Length distributions of detected transposable elements in the GRCh38 and JG1 genomes. Shown are *Alu*, SVA, and LINE1. Transposable elements and their subclasses were identified using RepeatMasker software (ver. 4.0.7) with the ‘- species human’ option. The resulting OUT format files were converted to BED format using the rmsk2bed command of BEDOPS software^59^ (ver. 2.4.35). Transposable elements disrupted by other elements were counted as distinct.

