## Supplementary Materials for "Construction and Integration of Three *De Novo* Japanese Human Genome Assemblies toward a Population-Specific Reference"

### SUPPLEMENTARY FIGURE LEGENDS

**Supplementary Fig. 1.** Karyotypes of the three subjects: **(a)** jg1a, **(b)** jg1b, and **(c)** jg1c. The arrow in panel **(a)** indicates the normal variation inv(9)(p12q13).

**Supplementary Fig. 2.** Harr plot of the alignment between chromosome 9 of GRCh38 and two largest scaffolds aligned to chromosome 9 from the **(a)** jg1a, **(b)** jg1b, and **(c)** jg1c assemblies, indicating that the two individual genomes harbor a possible shared inversion. 'Super-scaffold' is the default prefix designated by BionanoSolve software.

**Supplementary Fig. 3.** Workflow of the construction of JG1. **(a)** Workflow of the construction of each draft assembly. **(b)** Workflow of the integration of three draft assemblies. Rectangles indicate substrates such as reads, contigs, and scaffolds. Rectangles with rounded corners indicate software or processes.

**Supplementary Fig. 4:** Histogram of PacBio subreads length for **(a)** jg1a, **(b)** jg1b, and **(c)** jg1c. The length of each subread was calculated using the SAMtools (ver. 1.8) faidx command.

**Supplementary Fig. 5:** Histogram of Bionano data for **(a)** Nt.BspQI of jg1a, **(b)** Nb.BssSI of jg1a, **(c)** jg1b, and **(d)** jg1c. The length of each molecule was extracted from the BNX file.

**Supplementary Fig. 6:** Majority decision. **(a)** Schematic representation of majority decision approach. **(b)** Venn diagram of SNVs detected in JG1, jg1a, jg1b, and jg1c by comparison with hs37d5. The intersection relationship was inferred using the BCFtools (ver. 1.8) isec command.

**Supplementary Fig. 7:** Length distributions of detected transposable elements in the GRCh38 and JG1 genomes. Shown are *Alu*, SVA, and LINE1. Transposable elements and their subclasses were identified using RepeatMasker software (ver. 4.0.7) with the '-species human' option. The resulting OUT format files were converted to BED format using the rmsk2bed command of BEDOPS software[59] (ver. 2.4.35). Transposable elements disrupted by other elements were counted as distinct.

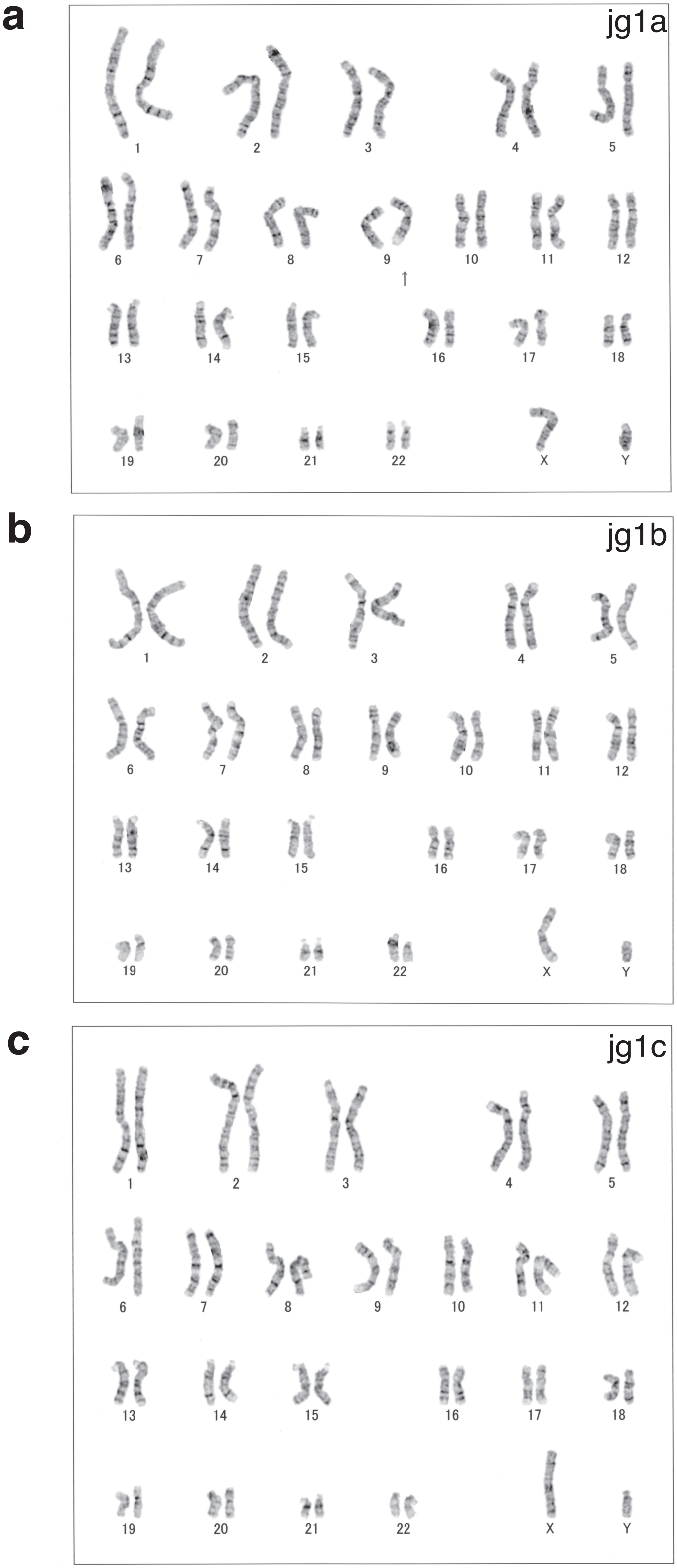


Supplementary Fig. 1; Takayama et al.

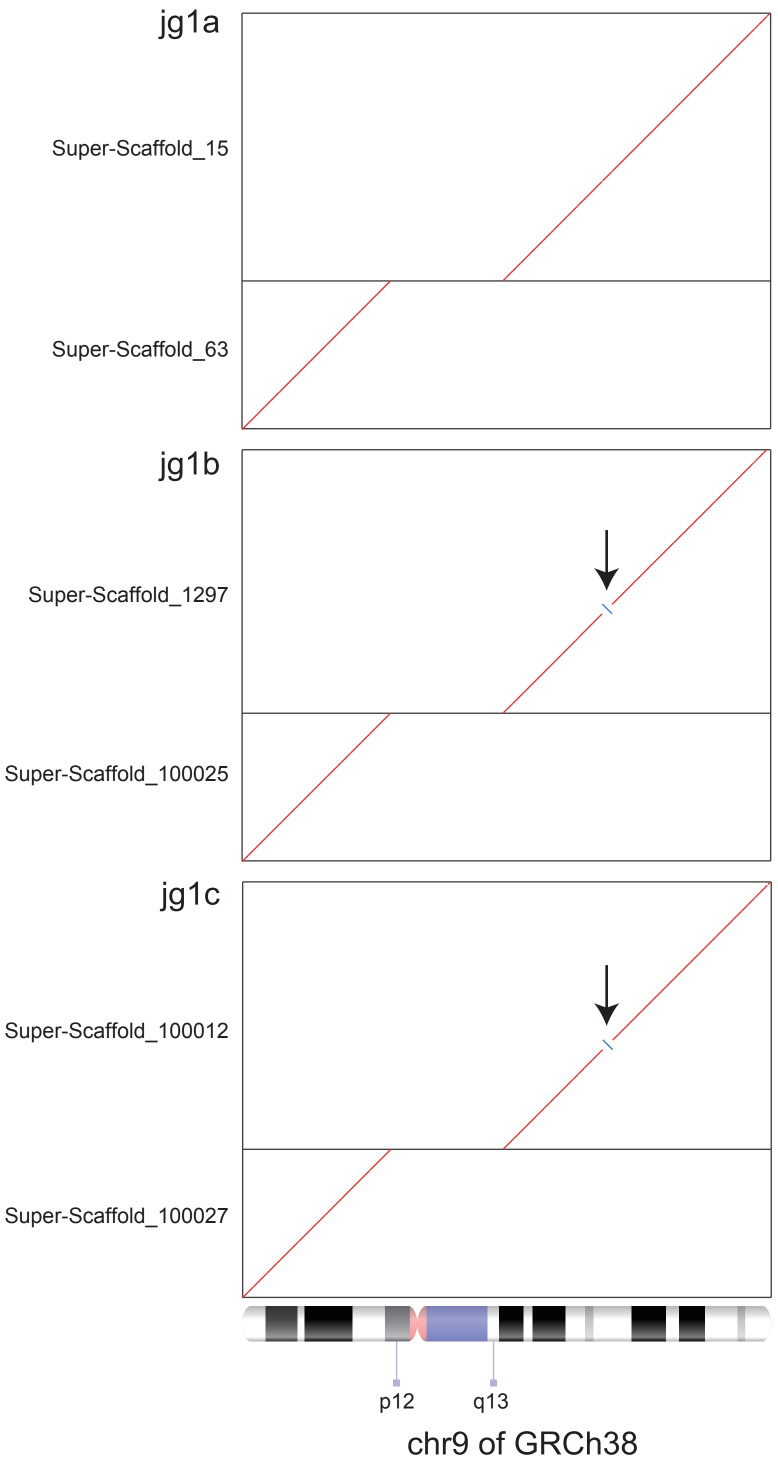


Supplementary Fig. 2

Takayama et al.

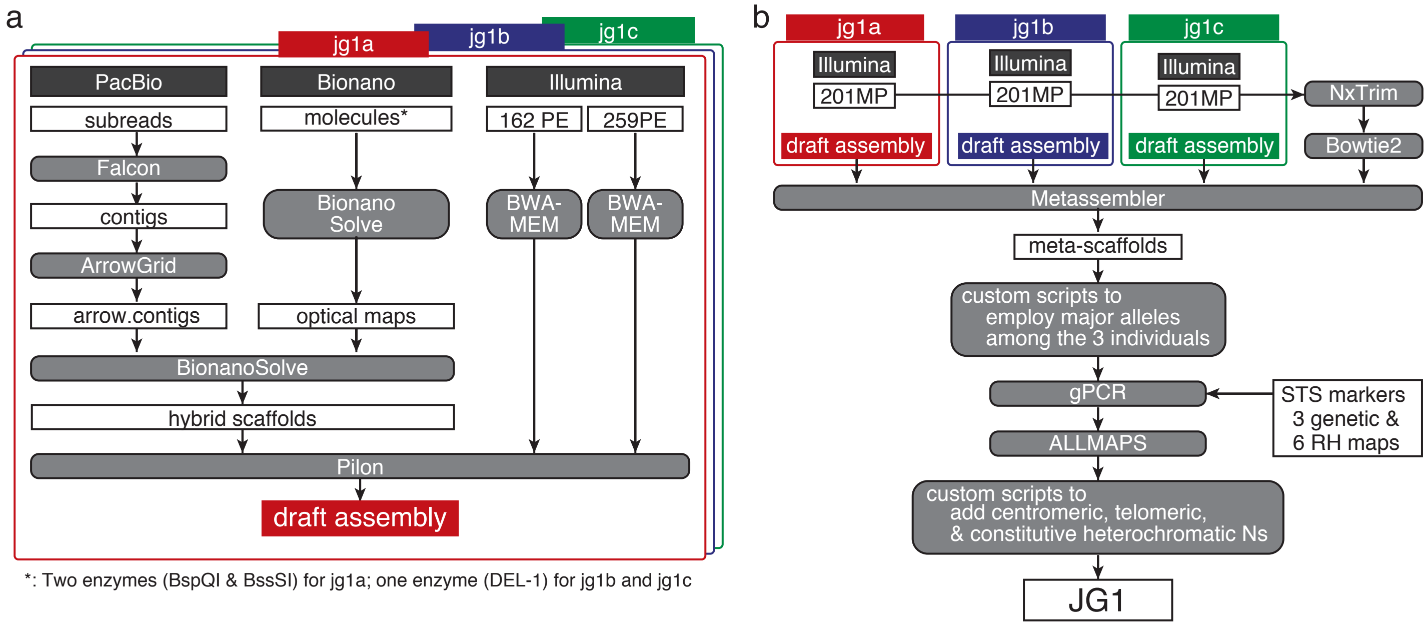


*: Two enzymes (BspQI & BssSI) for jg1a; one enzyme (DEL-1) for jg1b and jg1c

Supplementary Fig. 3; Takayama et al

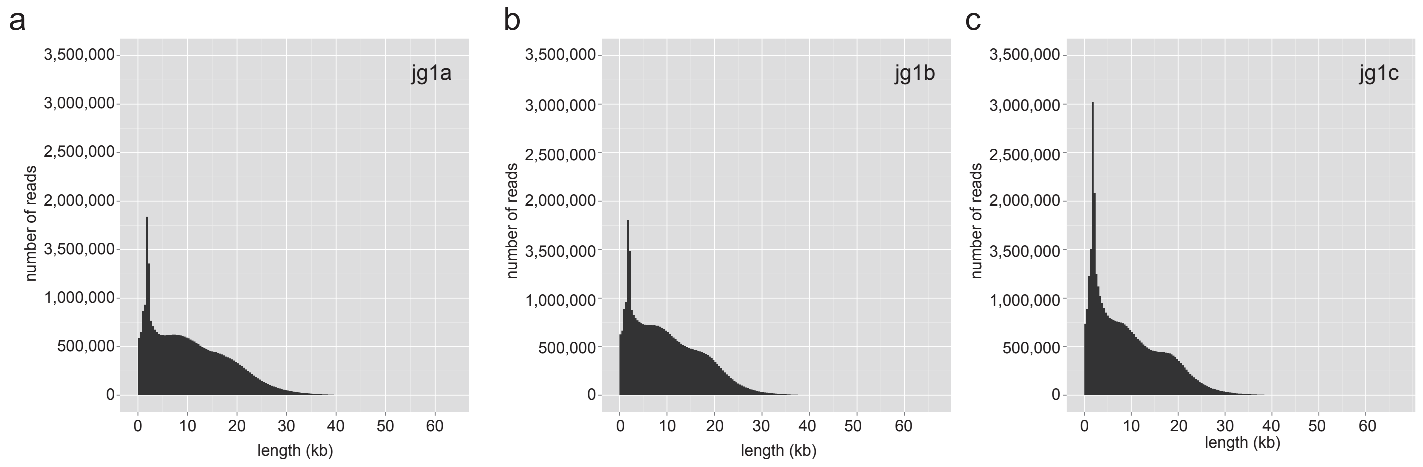


Supplementary Fig. 4

Takayama et al.

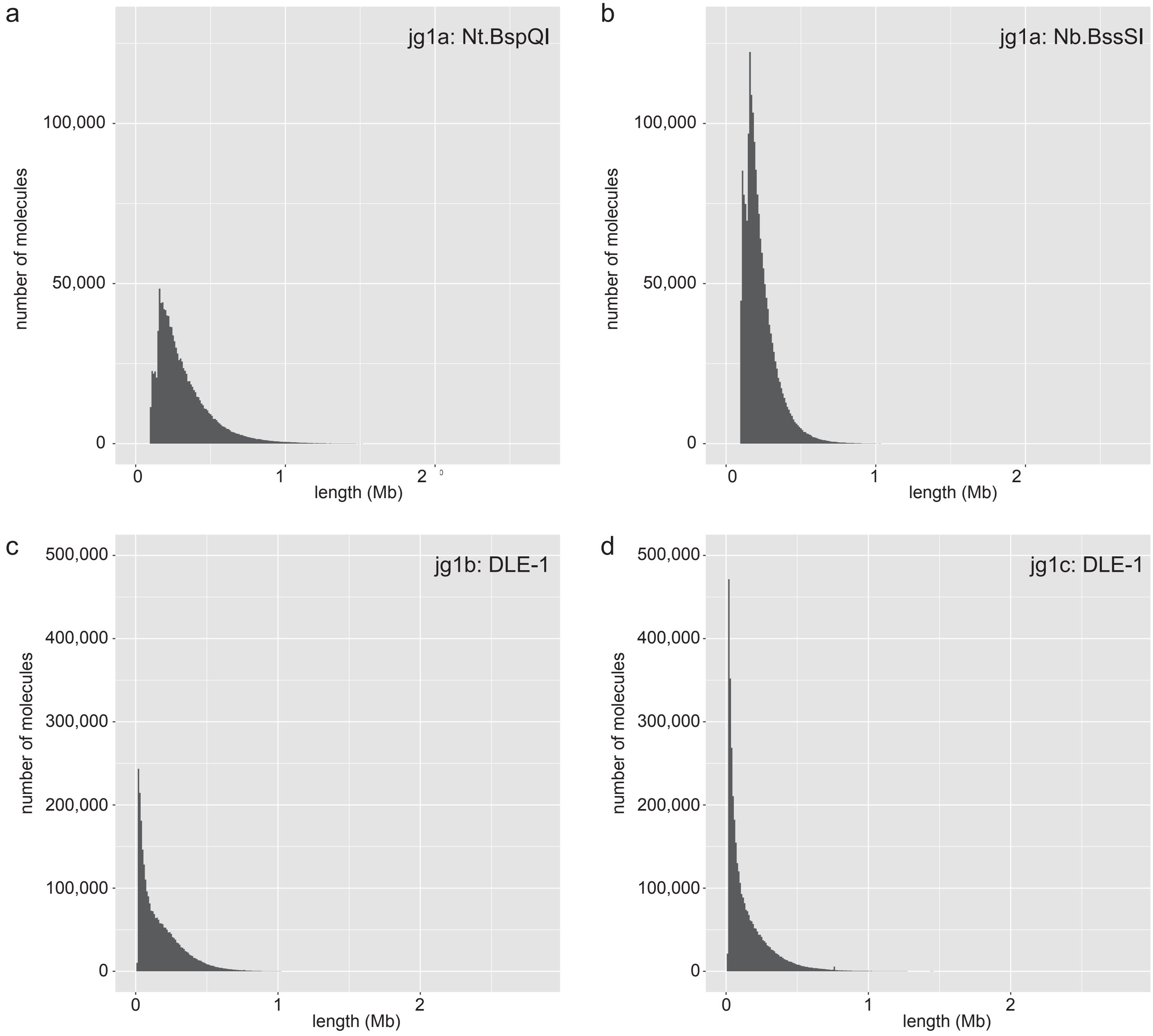


Supplementary Fig. 5

Takayama et al.

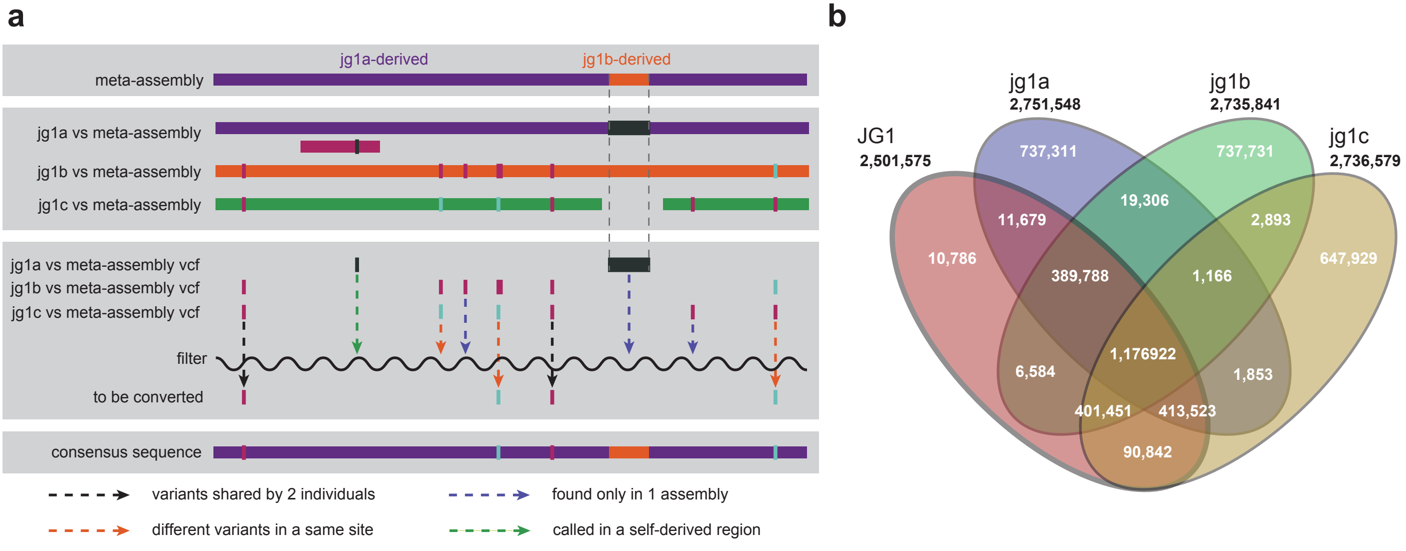


Supplementary Fig. 6

Takayama et al.

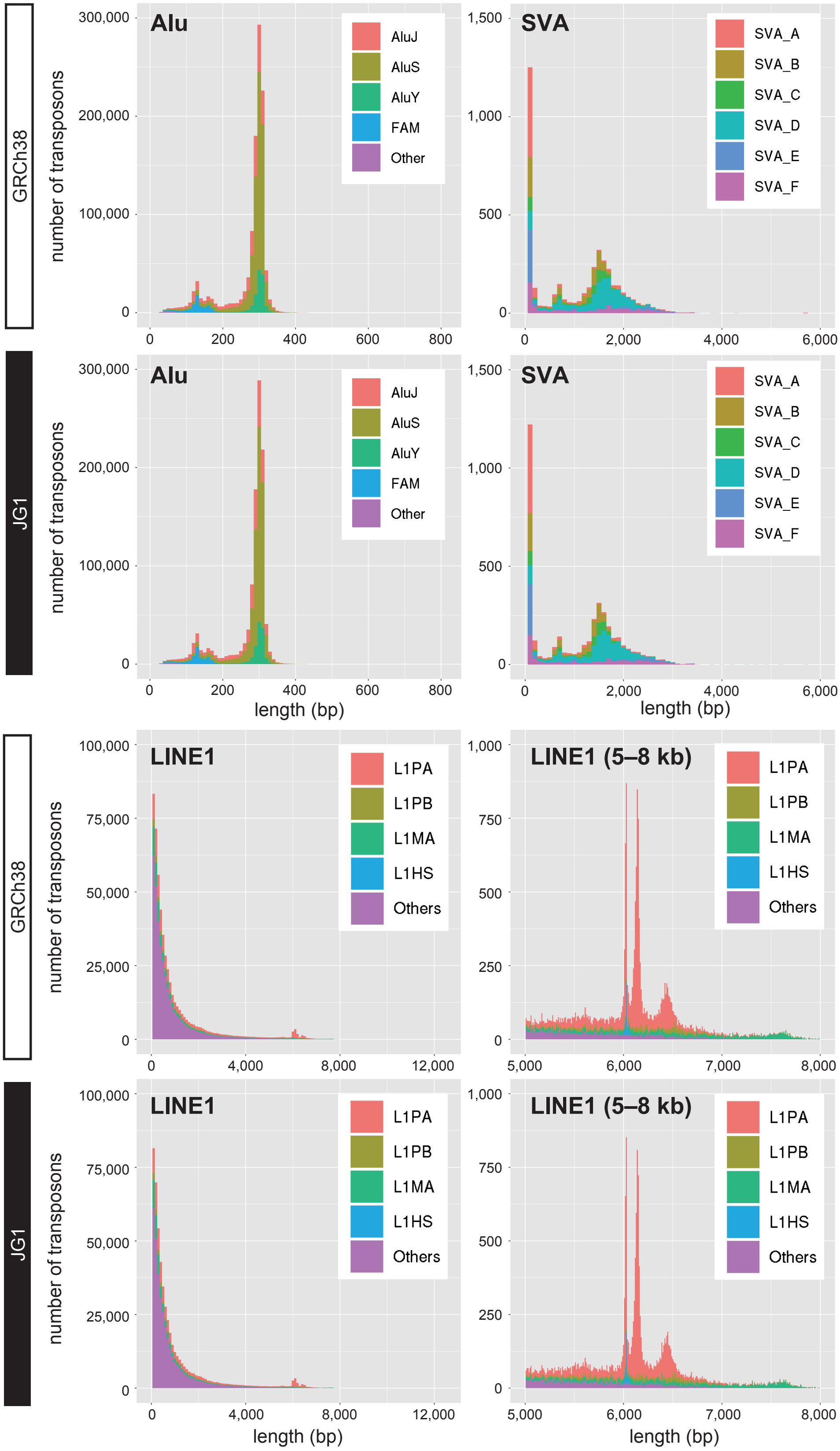


Supplementary Fig. 7

Takayama et al.

**Supplementary Table 1**. Basic statistics of PacBio subreads.

| Individual | Number of subreads | Sum of subread length (bp) | depth* |
|---|---|---|---|
| **jg1a** | 34,445,474 | 364,777,563,591 | 122X |
| **jg1b** | 36,798,731 | 370,437,373,175 | 123X |
| **jg1c** | 41,535,337 | 383,220,406,482 | 128X |

* Depth is calculated by assuming the genome size = 3.0 Gb.

**Supplementary Table 2.** Basic statistics of Bionano molecules.

| Individual | Enzyme | Number of molecules | Sum of molecule length (bp) | depth* |
|---|---|---|---|---|
| **jg1a** | BspQI | 1,156,682 | 368,075,072,000 | 123X |
| | BssSI | 1,834,771 | 418,513,858,000 | 140X |
| **jg1b** | DLE-1 | 2,840,733 | 480,476,071,000 | 160X |
| **jg1c** | DLE-1 | 3,594,225 | 524,851,027,000 | 175X |

* Depth is calculated by assuming the genome size = 3.0 Gb.

**Supplementary Table 3.** Basic statistics of Illumina paired-end and mate-pair reads.

| Method | Individual | read length (bp) | # of reads | Sum of read length (bp) | depth* |
|---|---|---|---|---|---|
| **paired end** | jg1a | 162 | 543,599,992 | 88,063,198,704 | 29X |
| | | 259 | 303,625,608 | 78,639,032,472 | 26X |
| | jg1b | 162 | 578,161,124 | 93,662,102,088 | 31X |
| | | 259 | 319,177,020 | 82,666,848,180 | 28X |
| | jg1c | 162 | 571,414,220 | 92,569,103,640 | 31X |
| | | 259 | 302,332,088 | 78,304,010,792 | 26X |
| **mate pair**** | jg1a | 201 | 189,189,310 | 38,027,051,310 | 13X |
| | jg1b | | 184,346,446 | 37,053,635,646 | 12X |
| | jg1c | | 185,928,504 | 37,371,629,304 | 12X |

* Depth is calculated by assuming the genome size = 3.0 Gb.

**all reads (before library separation)

**Supplementary Table 4.** Basic statistics of Bionano assembly.

| Individual | Enzyme | # of fragments | N50 (Mb) | Total length (Mb) |
|---|---|---|---|---|
| **jg1a** | BspQI | 4,761 | 1.179 | 3846.912 |
| **jg1a** | BssSI | 4,392 | 1.034 | 3202.036 |
| **jg1b** | DLE-1 | 581 | 41.761 | 3194.487 |
| **jg1c** | DLE-1 | 496 | 64.293 | 3481.086 |

**Supplementary Table 5.** Length of consecutive Ns inserted manually.

| chr | pter (bp) | cen (bp) | qter (bp) | References |
|---|---|---|---|---|
| **1** | 10,000 | 30,000,000 | 10,000 | 48–50 |
| **2** | 10,000 | 3,000,000 | 10,000 | |
| **3** | 10,000 | 3,000,000 | 10,000 | |
| **4** | 10,000 | 3,000,000 | 10,000 | |
| **5** | 10,000 | 3,000,000 | 10,000 | |
| **6** | 10,000 | 3,000,000 | 10,000 | |
| **7** | 10,000 | 3,000,000 | 10,000 | |
| **8** | 10,000 | - | 10,000 | |
| **9** | 10,000 | 30,000,000 | 10,000 | 48–50 |
| **10** | 10,000 | 3,000,000 | 10,000 | |
| **11** | 10,000 | - | 10,000 | |
| **12** | 10,000 | 3,000,000 | 10,000 | |
| **13** | 16,000,000 | - | 10,000 | 50 |
| **14** | 16,000,000 | - | 10,000 | 50 |
| **15** | 17,000,000 | - | 10,000 | 50 |
| **16** | 10,000 | 20,000,000 | 10,000 | 48–50 |
| **17** | 10,000 | 3,000,000 | 10,000 | |
| **18** | 10,000 | 3,000,000 | 10,000 | |
| **19** | 10,000 | 3,000,000 | 10,000 | |
| **20** | 10,000 | 3,000,000 | 10,000 | |
| **21** | 11,000,000 | - | 10,000 | |
| **22** | 13,000,000 | - | 10,000 | |
| **X** | 10,000 | 3,000,000 | 10,000 | |
| **Y** | 2,260,577 | 3,000,000 | 30,000,000 | 48, 51–53 |
